# LMD: Cluster-Independent Multiscale Marker Identification in Single-cell RNA-seq Data

**DOI:** 10.1101/2023.11.12.566780

**Authors:** Ruiqi Li, Rihao Qu, Fabio Parisi, Francesco Strino, Hainan Lam, Jay S. Stanley, Xiuyuan Cheng, Peggy Myung, Yuval Kluger

## Abstract

Identifying accurate cell markers in single-cell RNA-seq data is crucial for understanding cellular diversity and function. Localized Marker Detector (LMD) is a novel tool to identify “localized genes” - genes exclusively expressed in groups of highly similar cells - thereby characterizing cellular diversity in a multi-resolution and fine-grained manner. LMD constructs a cell-cell affinity graph, diffuses the gene expression value across the cell graph, and assigns a score to each gene based on its diffusion dynamics. LMD’s candidate markers can be grouped into functional gene modules, which accurately reflect cell types, subtypes, and other sources of variation such as cell cycle status.

We apply LMD to mouse bone marrow and hair follicle dermal condensate datasets, where LMD facilitates cross-sample comparisons, identifying shared and sample-specific gene signatures and novel cell populations without requiring batch effect correction or integration methods. Furthermore, we assessed the performance of LMD across nine single-cell RNA sequencing datasets, compared it with six other methods aimed at achieving similar objectives, and found that LMD outperforms the other methods evaluated.

## 1 Introduction

Single-cell RNA sequencing can currently reveal the transcriptomes of 10^4^ *−*10^7^ cells simultaneously [1]. These experimental approaches have enabled a better understanding of heterogeneity, cellular specialization and differentiation, and also provide insights into cell-cell signaling, spatial organization, and temporal dynamics leading to processes such as senescence, apoptosis, or oncogenesis [2]. An important task in single-cell data analysis is the identification of genes with high specificity to certain cell populations, which may indicate distinct cellular identities or states in this biological system.

Most marker identification methods [3–10] first cluster cells based on their similarity in the transcriptomic space and then detect genes by differential expression analysis. This approach has been criticized for its unreliable cluster assignments and inability to detect new cell subtypes defined by increased expression of specific genes [11–13]. Several state-of-the-art tools do not rely on an initial clustering step. For instance, SinglecellHaystack [11] and SEMITONES [13] utilize reference cells or grid points in the transcriptomic space to approximate cell geometry. They assess if a gene’s expression aligns with the arrangement of cells within this coordinate system. One limitation of SinglecellHaystack and SEMITONES is that they are confined to a specific range of length scales and may therefore fail to detect features at other length scales. Marcopolo[12] binarizes the expression level of a gene of interest and examines whether the cells expressing it are positioned near each other in a low-dimensional space. The proximity is defined by the average distance from each cell expressing the gene to their collective centroid. They assume that this proximity indicates the gene’s informativeness in terms of its association with a meaningful biological status. However, the binarization procedure can lead to a loss of quantitative information. Additionally, the positioning of the cell centroid and proximity assessments can be sensitive to outliers. Hotspot [14] employs a cell-cell similarity graph to identify informative genes with non-random expression patterns, using a modified Geary’s C. This ‘non-randomness’ is a general term that can include very different patterns: a gene being highly expressed in a certain region of the cell graph or showing a smooth expression gradient across the cell manifold. Although each pattern might suggest different biological interpretations, Hotspot does not provide an automated way to distinguish between these scenarios. This limitation, in turn, means it doesn’t impose constraints to ensure that the genes it reports are exclusively expressed in tight regions comprising highly similar cells. Consequently, the lack of specificity in some of the genes identified by Hotspot limits its effectiveness for clear and definitive cell type classification.

To identify the informative genes that categorize distinct cell subtypes or cellular states, we require that such a gene exhibit a “localized expression pattern” in the cell graph - the graph that represents cell-cell transcriptomic similarities. This is based on the assumption that cells of the same type and state will exhibit highly similar transcriptomic profiles, thereby aggregating in certain neighborhood(s) of the cell graph. To accomplish this goal, we propose a novel algorithm, Localized Marker Detector (LMD), designed to identify all localized gene markers at multi-scale resolutions. Specifically, LMD finds marker genes by (i) constructing a cell-cell affinity graph, (ii) diffusing the gene expression value across the cell graph, and (iii) assigning a score to each gene based on the dynamics of its diffusion process (Fig. 1A). Intuitively, the diffusion process takes longer to spread across the entire cell-cell graph when a gene is active in cells that are closer to each other on the cell-cell graph, or when the cells expressing the gene are a subset of a region that is less connected to the rest of the graph. The LMD’s markers can be used for various downstream applications. For instance, identifying gene modules based on co-localized genes can improve the characterization of cell groups by highlighting genes associated with specific cell types and states, providing deeper insights into their functions. Additionally, our approach allows for cross-sample comparisons to identify sample-specific or universally shared cell populations without the need for batch effect correction or integration methods, which often obscure important biological effects and lack theoretical justification (Fig. 1A).

**Fig. 1.**
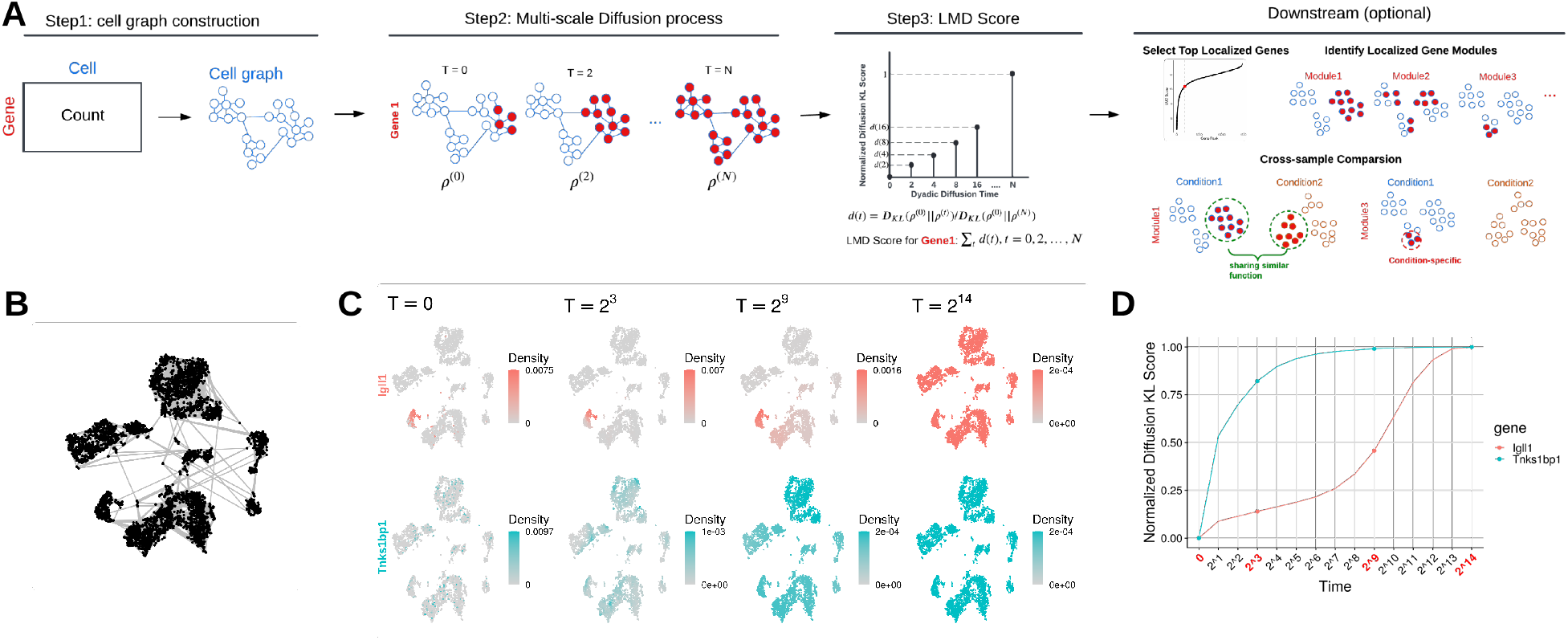
LMD Workflow: (A) Schematic illustration of the workflow of LMD (B) Construction of the cell kNN (k=5) graph. (C) Gene diffusion patterns (color-coded by density, computed as the initial or diffused state in each cell) for a localized gene *Igll1* (red) and a non-localized gene *Tnks1bp1* (cyan) overlaid on the t-SNE cell embedding at different time scales. (D) Normalized KL-divergence pattern of *Igll1* and *Tnks1bp1* during the diffusion process, with normalized KL-divergence against the time scale. Time scales in panel C are highlighted in red in panel D.

In this work, we benchmark LMD against several state-of-the-art marker identification algorithms on nine real single-cell datasets, demonstrating its ability to prioritize genes that are exclusively expressed in specific cell subpopulations, accurately identify known markers, and enhance the separation of cell groups with its top candidate markers. We then apply LMD to a mouse bone marrow dataset, grouping markers into distinct functional gene modules based on their co-localization in the cell graph; these modules delineate different cell states at multiple scales. Additionally, we used a mouse embryonic skin dataset to show that gene modules generated from LMD’s markers can be used to compare different conditions of the same system, identifying both condition-specific and universally shared cell populations and their associated genes. Our results highlight LMD’s ability to identify informative markers that enhance various downstream tasks, particularly the identification and characterization of functional cell groups.

## 2 Results

### 2.1 Overview of the LMD Method

Here, we outline the main three steps of LMD using a single cell RNA-seq mouse bone marrow dataset with FACS-based full length transcript technique from the Tabula Muris consortium[15] as an example (Fig. 1, S1A).

#### Cell graph construction

The first step of LMD involves computing a cell-cell affinity graph that shows which cells are sufficiently similar to each other based on a chosen similarity metric. We construct a k-nearest neighbor (kNN) graph where each cell is represented by a node, and two nodes are connected by an edge if either of their respective transcriptomes is among the closest *k* neighbors of the other according to a given metric. The metric can be any function that quantifies the similarity of cells, such as the Euclidean distance between their transcription profiles after reducing the dimensionality with Principal Component Analysis (PCA). The graph is undirected and unweighted, i.e., the connections are symmetric and all edges have equal weight. Fig. 1B shows a cell kNN (*k* = 5) graph constructed from the Euclidean distances between the transcription profiles after PCA.

#### Multi-scale diffusion process

In this step, we examine the gene expression pattern over the cell graph. We first normalize the expression of a gene into a probability distribution over cells, which we define as the initial state vector of the corresponding gene. Next, we apply a diffusion process with a series of time scales - a technique widely adopted in various fields [16–18] - and track how this state vector changes as a function of time. If the graph is connected, given enough time, all nodes will reach a non-zero value regardless of initial states, and the equilibrium state vector for our setup is a uniform distribution.

Some initial states are localized, meaning that most nodes in the graph have zero values except for the nodes in a specific subgraph (Fig. 1C, top panel). We observe that for a more localized initial state, it takes a longer time to reach a non-zero value at every node and then reach the equilibrium state (Fig. 1D). To quantify this effect, we measure changes in the state vector as a function of time by computing the Kullback-Leibler divergence (KL-divergence) between the state at time *t* and the initial state at time *t*_0_. To adjust for the effect where initial states distributed across different numbers of nodes result in varying convergent values, we normalize this time-dependent divergence by the KL-divergence between the initial state and the equilibrium state, and term this quantity as normalized KL-divergence. As expected we observe that the normalized KL-divergence for more localized initial states increases at a slower rate during the early stages of the diffusion process, indicating that the redistribution of values primarily occurs between nodes with similar values. In contrast, for initial states that are not localized, the diffusion process quickly redistributes values from higher-value nodes to neighboring lower-value nodes at the beginning. This results in a rapid increase in normalized KL-divergence.

#### The LMD-score (LMDS)

To summarize the time-dependent profile of the normalized KL-divergence, we calculate the area under the curve (Fig. 1D), which we term as the LMD-score (LMDS). A lower LMDS indicates a more localized initial state. In the context of gene expression profiles, we find that genes expressed in specific regions of the cell graph are assigned a lower LMDS, implying LMDS can distinguish localized genes from the genes non-specific to any local regions of the cell graph. This is the main idea of LMD.

### 2.2 LMD prioritizes localized markers

To better understand the difference between LMD and the other state-of-art methods – Seurat v4 default, Hotspot, SEMITONES, Marcopolo, and singleCellHaystack – we compared their gene rankings on the Tabula Muris bone marrow FACS dataset. Among the methods, Seurat v4 showed the highest concordance with LMD, while singleCellHaystack had the lowest (Fig. 2A). Candidate markers selected by LMD were typically expressed in one or several compact cell neighborhoods, whereas other methods either prioritized markers lacking locality or overlooked markers expressed in a subset of a cluster (Fig. 2B). In detail, Seurat v4 identifies genes through differential expression and Hotspot prioritizes genes with global variation, thus both methods may overlook the confinement of markers to specific cell groups where their candidate genes can be expressed at high levels in a cluster but also at lower levels in dispersed cells outside the cluster; Marcopolo’s markers often correspond to a noisy and less localized signal; SEMITONES and SingleCellHaystack missed markers expressed in cell subsets smaller than their pre-defined fixed length scale.

**Fig. 2.**
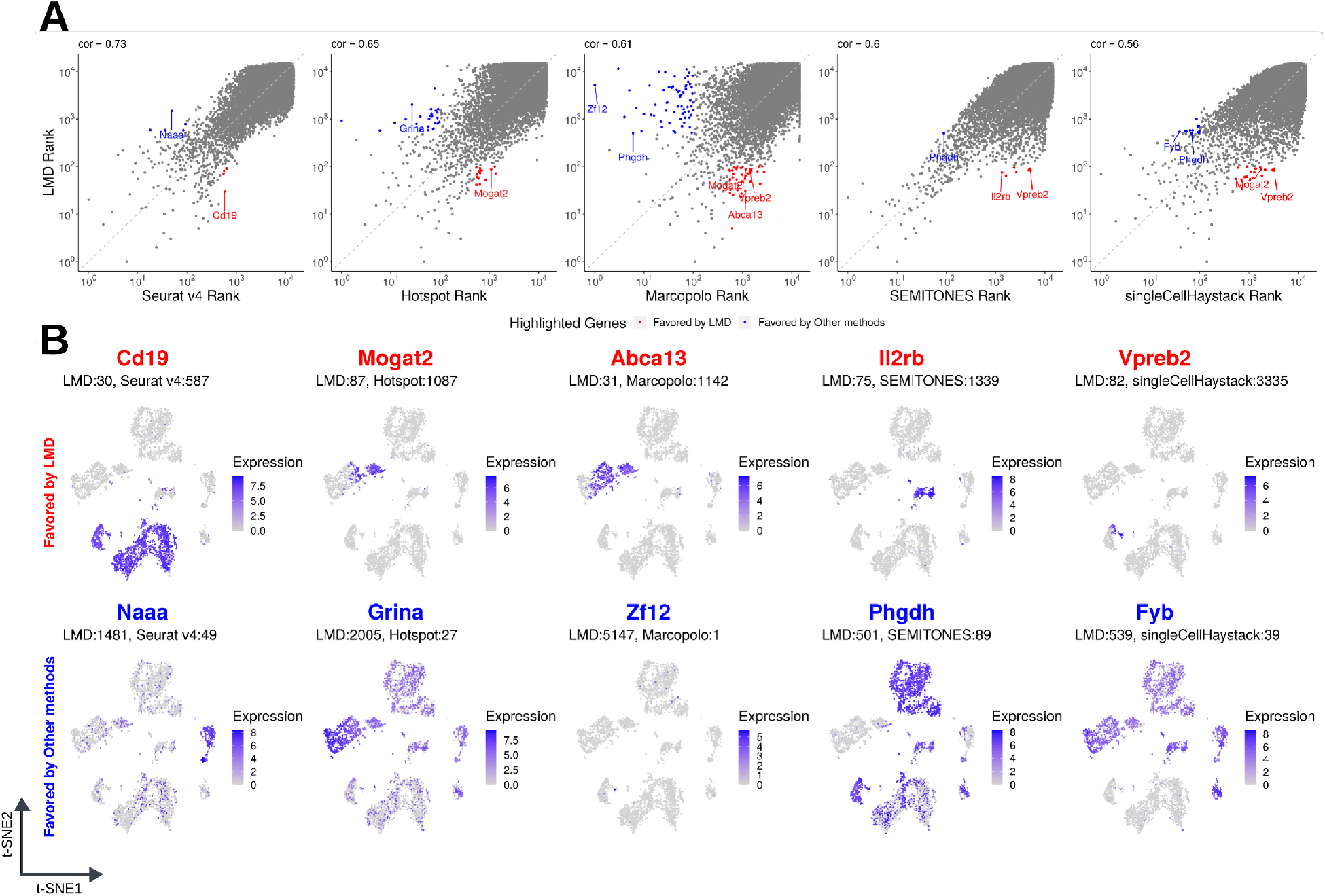
Gene Ranking Comparisons between LMD and Other Methods. (A) Comparison of gene rankings by LMD against Seurat v4 default, Hotspot, Marcopolo, SEMITONES, and SingleCellHaystack, with concordance measured using the Pearson correlation coefficient of log-transformed rankings (base 10). Highlighted genes fall within LMD’s top 100 ranked genes but are outside the top 500 ranked genes of the other method (red), and vice versa (blue). (B) Top localized genes selected by LMD but not highly ranked by other methods and vice versa. The rankings of these genes across all methods can be found in Table S1.

### 2.3 LMD identifies known markers with high accuracy

We evaluated LMD on its ability to identify known markers using nine real scRNA-seq datasets: three datasets from the Tabula Muris consortium[15] and six datasets from the Azimuth demo datasets[19]. We compared LMD performance to a suite of marker gene identification algorithms, including singleCellHaystack[11], Hotspot[14], SEMITONES[13], MarcoPolo[12], standard DEG workflow based on clustering and Wilcoxon rank sum test from Seurat v4[3] and Highly Variable Gene (HVG) implemented in Seurat v4 with VST option. To validate performance, we used two distinct true marker sets as suggested in [12]. The first set was derived from marker databases, and the second set consisted of the top 100 genes with the largest expression fold change across cell types (see Section. 4.3). We measured the AUROC of the methods for each dataset and found that LMD outperforms all other methods in most cases (Fig.3). For the first true marker set, LMD ranked first in 5 out of 9 datasets, and for the second true marker set, it also ranked first in 5 out of 9 datasets. Overall, LMD achieved the best performance for both true marker sets based on the median rank across all datasets.

**Fig. 3.**
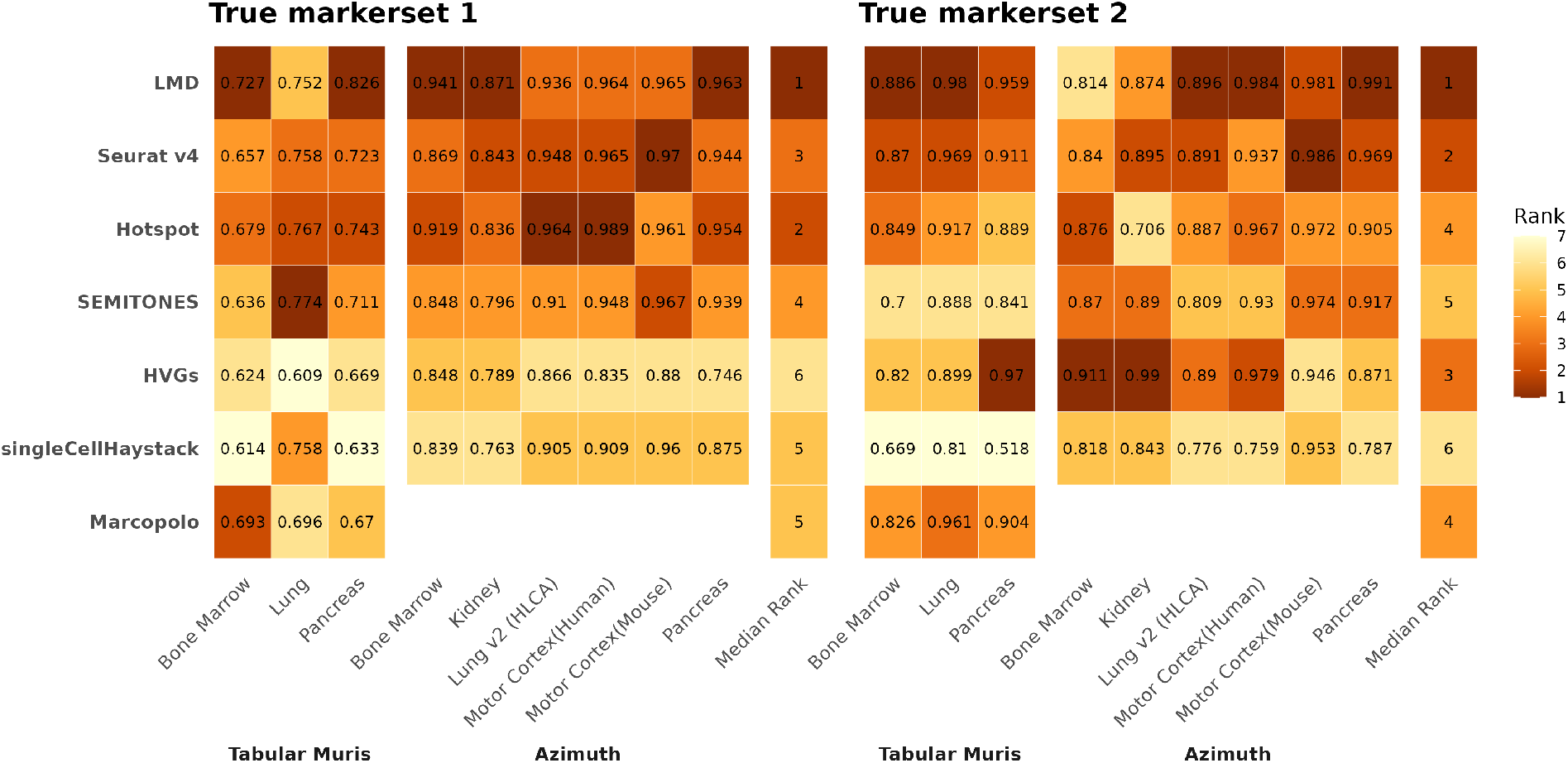
Comparison of AUROC for identifying cell type markers across 7 methods using 9 real datasets (Tabula Muris and Azimuth demo datasets). Each block represents the AUROC score for a method on a dataset, with the number showing the AUROC value and the color indicating the method’s rank. Marcopolo (CPU) was only run on the Tabula Muris dataset due to runtime and hardware restrictions. Two true marker sets: 1 – genes listed in marker databases (CellMarkerDB and Azimuth marker list); 2 – the top 100 genes with the maximum log fold change across pre-annotated cell types.

We also compared the ability of markers from each method to enhance separability between cell groups on the Tabula Muris bone marrow FACS dataset using the density index[20]. The density index measures ‘clumpiness’ in cell distribution by comparing the average closeness of nearest neighbors to the average distance between random cell pairs. A higher value indicates a more compact and clumped cell distribution. LMD achieved a higher density index than other methods (Fig. S2A). The density index for LMD peaked around the top 300 genes, suggesting that even a small number of LMD genes is informative for separating diverse cell populations. Furthermore, when focusing on the granulocyte subset of this dataset, LMD still maintained a higher or comparable density index, demonstrating its ability to identify informative markers that help distinguish cell subpopulations, even in more homogeneous cases (Fig. S2B).

LMD also demonstrated reasonable runtime performance (Section 4.3, Table S2). For example, it processed 1,564 cells in approximately 30 seconds and 30,651 cells in about 44 minutes.

### 2.4 LMD markers unveil functional modules and cell states across varied length scales in mouse bone marrow

#### Gene modules

We demonstrate the flexibility of LMD markers in characterizing functional cell groups using the Tabula Muris bone marrow FACS dataset, ranging from broad gene expression across multiple regions to fine-grained expression within a single cluster. First, we selected the top 1,741 LMD candidate markers based on the knee point of the gene LMDS curve. To characterize various expression patterns, we grouped these markers into 34 distinct gene modules (Section 4.2, Fig. S3A and S3B): of these, 20 were enriched for at least one GO molecular function term (Benjamini-Hochberg adjusted p-value < 0.05), and 24 were enriched for at least one pathway from the Reactome Pathway Database[21] (Benjamini-Hochberg adjusted p-value < 0.05). For detailed information on each module, see Section S3.1. As expected, given the locality of the underlying markers, each module was associated with specific regions of the cell-cell affinity graph, which enables the characterization of the data at different length scales (Fig. S3C). At a broader length scale, we found several modules that were jointly expressed across multiple local regions, for instance, modules associated with progression through the cell cycle. At cell-type resolution, LMD identified several modules that aligned well with known annotations. Finally, at a finer length scale, LMD revealed several gene modules, each associated with a specific subset in a pre-annotated cell type, suggesting the potential of using LMD to define novel cell types at a high resolution. Moreover, inspecting certain gene modules may elucidate the different cell states along a dynamic biological process (e.g., B lymphocyte differentiation) and facilitate the discovery of novel cell subtypes in this system.

#### Broad length-scale, larger data structures

LMD identified three gene modules significantly enriched for cell cycle phases based on Reactome pathway analysis. Specifically, modules 22, 20, and 19 were significantly enriched for the S phase, S/M phase, and M phase, respectively (Benjamini-Hochberg adjusted p-value < 0.05, Fig. 4A). We confirmed this result using an alternative Tabula Muris bone marrow dataset with microfluidic droplet-based 3’-end counting technique [15] (Fig. S1B). In both datasets (bone marrow FACS and droplet), the depiction of these gene modules on the cell t-SNE embedding closely matches cell annotations derived from Seurat cell cycle scores, which rely on established cell cycle markers (Fig. 4B). The modules derived from LMD show incremental activation of markers, revealing the dynamics of cell-cycle progression (Fig. 4B and S5).

**Fig. 4.**
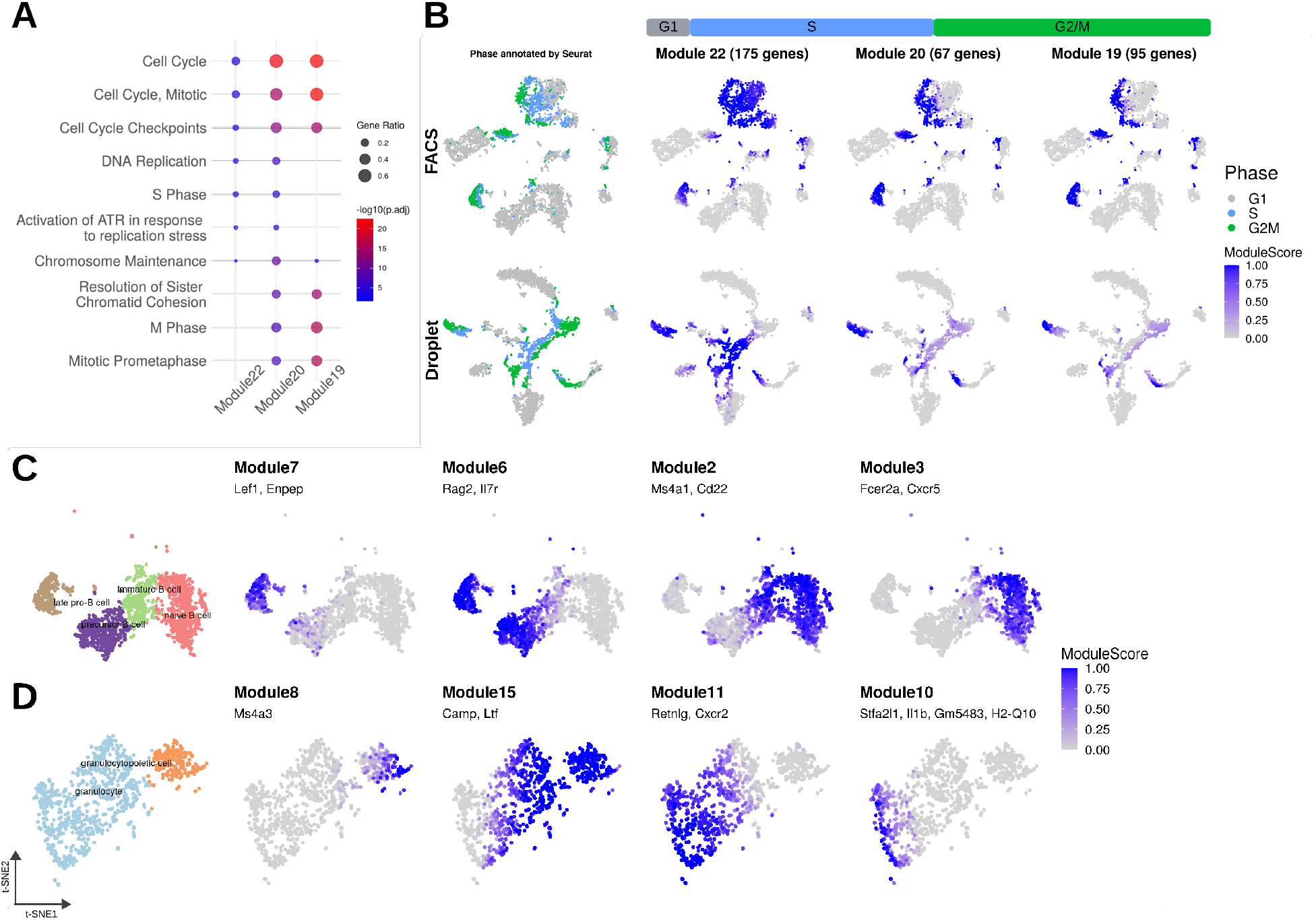
Gene module specificity in cell cycle stages and within cell subsets. (A-B) Gene modules associated with the cell cycle. Gene modules were identified from the Tabula Muris bone marrow FACS dataset and their function was classified by Reactome pathway analysis. Modules 22, 20, and 19 were associated with S phase, S/G2/M and G2/M phases, respectively. (A) Bubble plot of the top 5 significantly enriched Reactome pathways (Benjamini-Hochberg adjusted p-value < 0.05) for each module. Bubble color indicates log(adjusted p-value, base 10), and the size represents the fraction of module genes associated with the pathway. (B) Module scores in two Tabula Muris bone marrow datasets, and reference cell-cycle annotation from Seurat (left). (C) Gene modules associated with B cells at different stages of maturation. Module 7 is associated with pro-B cells, module 6 with pre-pro-B cell, module 2 with immature and mature B cells, and module 3 with naive B cells. (D) Gene modules associated with the production of granulocytes. Module 8 contains genes expressed during granulocytopoiesis, module 15 contains genes common for granulocytopoietic cells and granulocytes, module 11 contains genes involved in granulocyte maturation, and module 10 may represent a rare granulocyte subgroup.

#### Cell-type resolution

LMD identified several modules that demonstrated specificity to pre-annotated cell types. The strength of this association was quantified using the Jaccard Index (see Section 4.2). Due to limited cell numbers, we merged T cell subtypes into ‘T cell’ and NK cell subtypes into ‘NK cell’. Of the resulting 18 cell types, 11 contained at least one distinctly activated module (Fig. S4A and S4B). The relationship between these modules and cell types was further validated as the modules incorporated canonical cell type markers used for annotating this Tabula Muris bone marrow dataset (see legends of Fig. S4). We performed cross-dataset verification on the droplet dataset and found that modules identified as specifically activated in particular cell types in the FACS dataset also showed high specificity for the same cell types in the droplet dataset (Fig. S4C).

#### High resolution, beyond cell types

LMD identified gene modules that trace the B lymphocyte differentiation process (Fig. 4C) or various developmental stages and subtypes of granulocytes (Fig. 4D). Modules associated with B lymphocytes are substantiated by the inclusion of genes with well-established roles in this differentiation process: module 7 (*Lef1* and *Enpep* (also known as *BP-1*)) distinguish pro-B cells[22, 23]; module 6 (*Rag2* and *Il7r*) represents pre-pro-B cell stage [24, 25]; module 2 (*Ms4a1* (also known as *Cd20*) and *Cd22*) pertains to B cell activation and regulation, marking both immature and mature B cells in the bone marrow [26, 27]; module 3 (*Fcer2a* (also known as *Cd23*) and *Cxcr5*) characterizes naive B cells [28, 29]. We similarly identified at least 4 modules associated with granulopoiesis: module 8, exclusively expressed in granulopoietic cells, including *Ms4a3*, a marker gene used to label bone marrow granulocyte-monocyte progenitors (GMP) and GMP progenies [30]; module 15, featuring secondary granules genes *Camp* and *Ltf*, expressed in both granulocyte and granulopoietic cells [31]; module 11, specific to a subset of granulocytes and containing *Retnlg* and *Cxcr2*, which suggests a potential role in neutrophil mature and release [32, 33]; lastly, module 10, characterized by genes implicated in pyroptosis and antigen processing such as *Stfa2l1, Il1b, Gm5483*, and *H2-Q10* [34–36], which may represent a rare granulocyte subgroup engaged in the immune response against intracellular pathogens.

### 2.5 LMD characterizes shared and mutant-specific cell populations in mouse dermal condensate genesis

Mouse hair follicle dermal condensates (DCs) appear in the dermis of the skin around embryonic day 14.5 (E14.5) and play a critical role in hair follicle development[37]. The differentiation of DC cells is influenced by the cooperative actions of Wnt/*β*-catenin and sonic hedgehog (SHH) signaling pathways. In wild type (WT) mice, Wnt signaling is activated prior to E14.5 as a gradient spanning the upper dermis, while SHH is expressed by hair follicle epithelial cells around E14.5, inducing SHH activation in the Wnt-active dermal cells beneath the hair follicle epithelium. Together, they regulate the emergence and differentiation of dermal condensates[37–39]. The SmoM2YFP mutant (SmoM2) induces uniformly high SHH activation across the dermis, resulting in the early appearance of DC-like structures at E13.5[37].

We collected mouse skin samples from E13.5 and E14.5 WT and paired SmoM2 mutant embryos for 10X scRNA-seq. We used LMD to examine the functional cell groups that are conserved between SmoM2 and WT, as well as those that reflect SmoM2-specific changes. We focused on the E13.5 SmoM2 sample and identified 17 gene modules (Fig. S8). Among them, we identified modules that: 1) reveal consistent fine-grained cell cycle phases across SmoM2 and WT (Fig. 5B), 2) reflect different cell stages in DC genesis of SmoM2 mutant (Fig. 5C), and 3) a novel module that highlights a special cell subset with a consistent transcriptomic profile in both E13.5 and E14.5 mutants but is absent in the WT condition (Fig. 6A and S14).

**Fig. 5.**
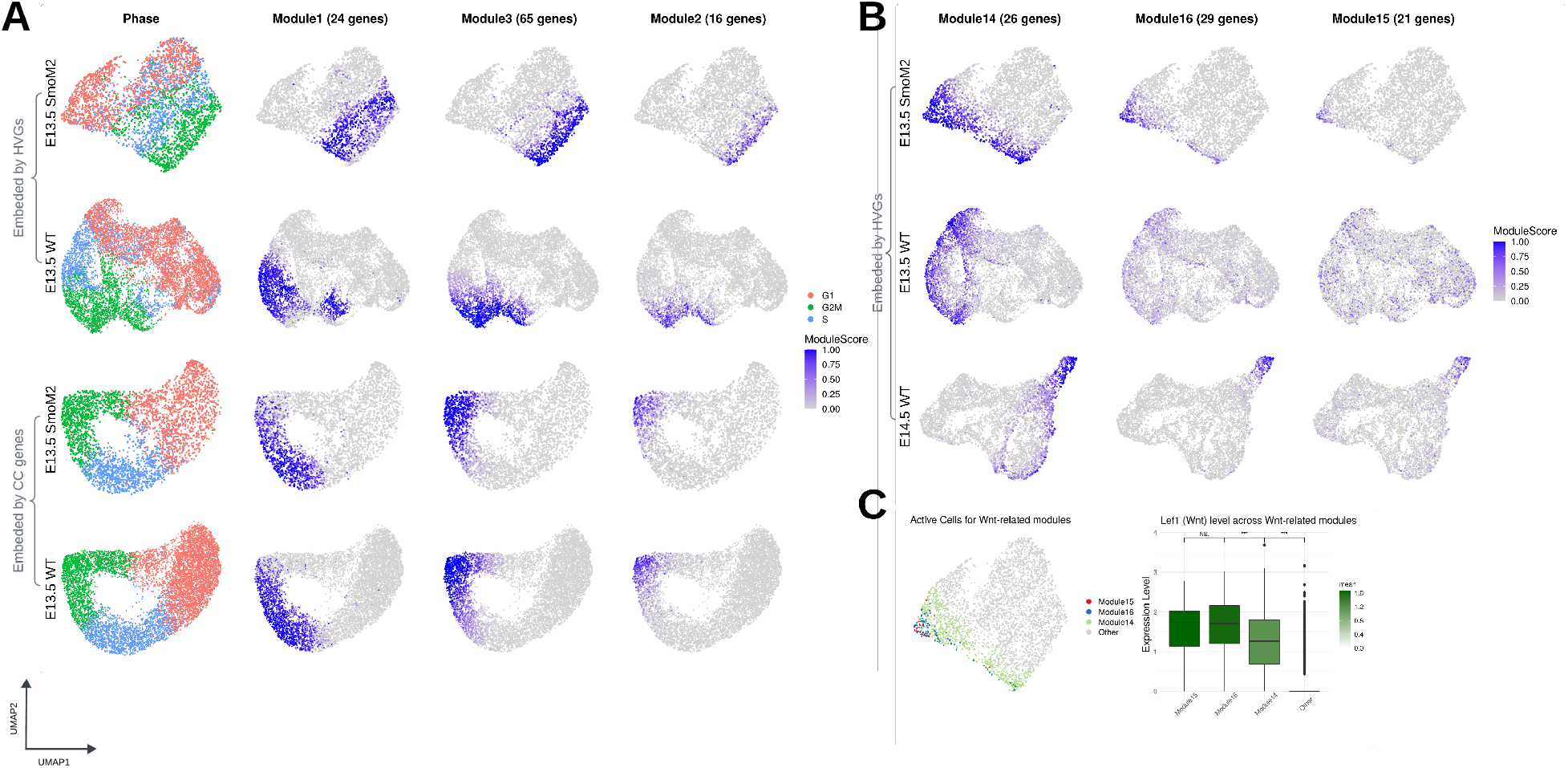
Comparative analysis of gene modules between samples in DC genesis. (A) Module scores of three cell cycle-associated gene modules on E13.5 SmoM2 and paired WT, with cell cycle annotations obtained from Seurat for reference (left). Different UMAP embeddings: by top 2000 HVGs of E13.5 SmoM2 (first row), top 2000 HVGs of E13.5 WT (second row), and merged embeddings of E13.5 SmoM2 and E13.5 WT by Seurat cell cycle genes (third row: SmoM2; fourth row: WT). (B) Module scores of three Wnt-associated gene modules on E13.5 SmoM2 (first row), paired E13.5 WT (second row) and E14.5 WT (third row). (C) Left: Cells categorized by three Wnt-associated gene modules (module score > 0.5, mutually exclusive: cells assigned to innermost module). Right: *Lef1* (Wnt) level in three modules: The box represents the interquartile range (IQR), with the line inside the box indicating the median. Whiskers extend to a maximum of 1.5×IQR beyond the box, with outliers represented as individual points. A Wilcoxon test was performed between adjacent boxes, with “NS” for *p* > 0.05 and “***” for *p* < 0.001.

**Fig. 6.**
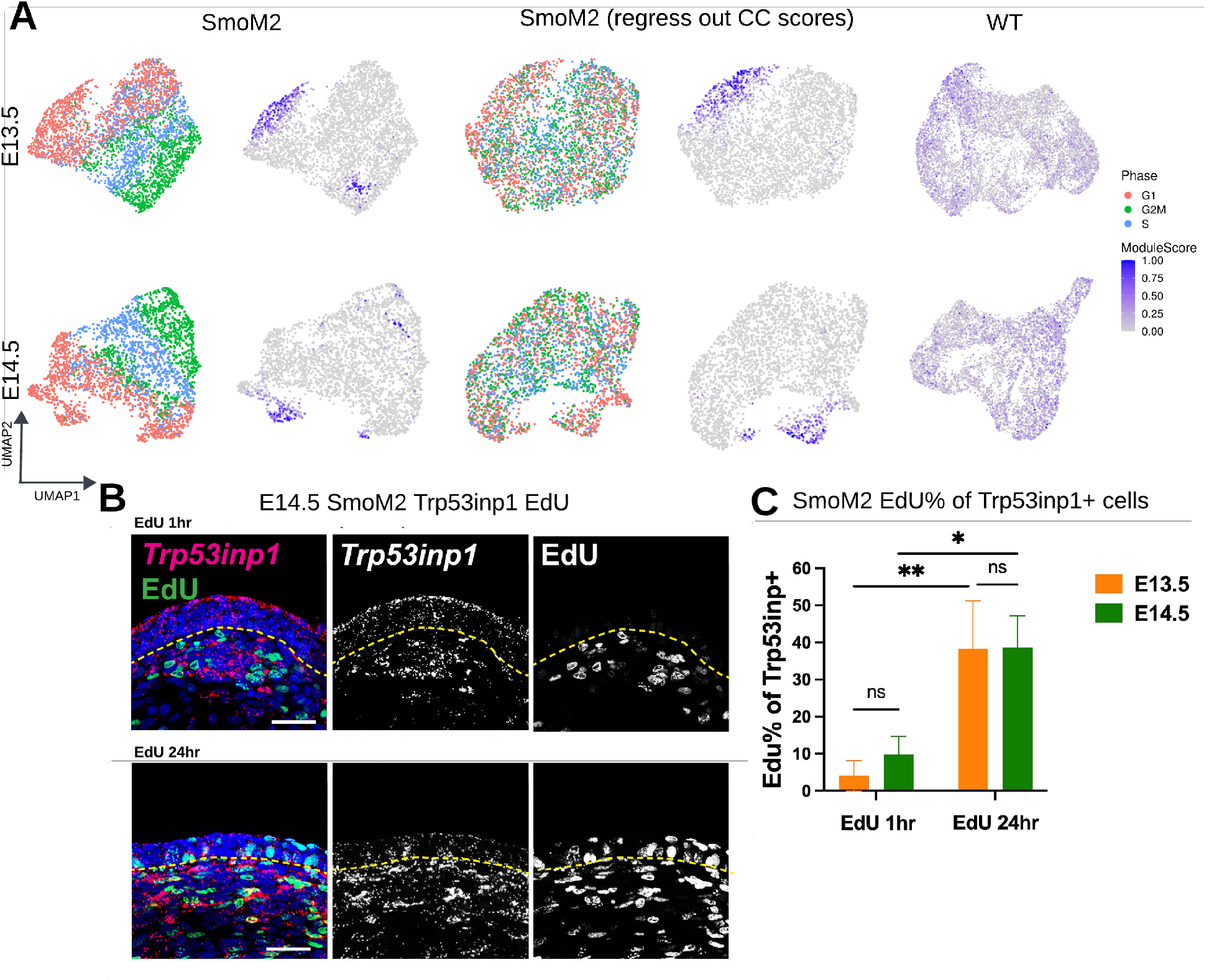
A gene module capturing mutant-specific cell subsets. (A) Module scores of module 12 on E13.5 SmoM2 (first row, columns 2 and 4) and paired WT (first row, column 5), and E14.5 SmoM2 (second row, columns 2 and 4) and paired WT (second row, column 5), with cell cycle annotations from Seurat (columns 1 and 3). Activated cells (module score > 0.5) in E13.5 SmoM2: 114 G1, 34 G2M, 9 S cells; in E14.5 SmoM2: 44 G1, 16 G2M, 4 S cells. UMAP embeddings: standard scaling (columns 1 and 2) and scaling regressing out cell cycle scores (columns 3 and 4). (B) FISH images (scale bar = 50 *µm*) showing the spatial distribution of *Trp53inp1* in the upper dermis of SmoM2 pulsed with EdU at E14.5 (first row) and 24 hours prior at E13.5 (second row). EdU is a nucleotide that is incorporated by cells in the S phase. (C) %EdU+ of *Trp53inp1* + cells at E13.5 and E14.5, pulsed with EdU (EdU 1hr) and 24 hours prior (EdU 24hr). Data as mean*±*SEM, one-way ANOVA with Tukey’s HSD test was performed between groups, with “NS” for *p* > 0.05, “*” for *p* < 0.05, “**” for *p* < 0.01.

#### LMD reveals functional similar cell populations in mutant and wild-type samples

We identified three distinct gene modules that capture fine-grained phases along the cell cycle procession: module 1 is characterized by genes typical of the S to G2/M transition, including *Hist1h2ap, Hist2h2bb, Hist1h2ae*[40]; module 3 exhibits genes expressed in the G2/M phase, such as *Cenpf, Cenpe* and *Cdk1* [41–43]; while module 2 is enriched for genes prevalent during late G2 and M phases, such as *Nek2, Kif14*, and *Kif18b* [44, 42] (Fig. 5A, first row). We also observed the localized pattern of these three modules in the paired E13.5 WT sample (Fig. 5A, second row). To investigate the functional connections of these localized patterns between SmoM2 and WT, we merged two samples and re-embedded cells by cell cycle marker genes (Section 4.3). Notably, despite the drastic transcriptome-wide differences between SmoM2 and WT, the new embedding generated using cell cycle marker genes aligns well between the two conditions (Fig. S7). In this new embedding, we noted that cells highlighted by these modules demonstrated good overlap between SmoM2 and WT, indicating the presence of these cell cycle phases across both samples.

The functional connections of localized patterns between two samples can also be validated by independent gene modules in each sample that share similar functions. Specifically, we apply LMD to each sample separately to identify gene modules. To pair gene modules between samples, we project the gene modules from one sample onto the other and match them based on their co-localization (Section 4.2). We then inspect whether the paired gene modules share similar biological functions through pathway analysis. For the three CC-associated modules identified from SmoM2, we projected them onto the WT sample and consistently found that each module co-localized with a corresponding module identified from the WT, with the minimum Jaccard index > 0.8 (Fig. S12A). Additionally, these paired modules are significantly enriched in common cell cycle-related pathways from the Molecular Signatures Database (MsigDB)[45] (Fig. S12B).

With the aforementioned points, LMD identifies conserved biological functions across different samples by projecting gene module expression patterns from one sample onto another. This approach detects localized patterns and corresponding cell subsets in the second sample without requiring cell integration or alignment between samples, eliminating reliance on “batch correction” methods that often fail to differentiate between batch effects and true biological variation.

#### LMD captures different cell states in the DC genesis mutant model

Previous work showed that higher levels of Wnt signaling correlate with increased DC formation in response to high SHH activation [37]. We identified three modules in E13.5 SmoM2–modules 14, 16, 15–that represent cell subsets with incremental activation of genes within the Wnt signaling gradient (Fig. 5). Module 14, which includes *Lef1*, a direct downstream target of Wnt/b-catenin signaling[46, 47], captures the broadest group of cells engaged in Wnt signaling. This is corroborated by its co-localization with two other modules from the E13.5 WT and E14.5 WT samples, with all three modules significantly enriched for the Wnt beta-catenin signaling pathway (Fig. S12). Module 15, containing the DC marker *Sox2* [48], captures the narrowest subset of cells representing DCs. It co-localizes with module 1 from the E14.5 WT sample, which also contains *Sox2* (Fig. S12). However, the localized pattern is absent in the E13.5 WT sample (Fig. 5B). This is consistent with the biological observation that early DC-like structures begin to form in the SmoM2 mutant at around E13.5, while DC emergence in WT happens around E14.5. Module 16 represents an intermediate stage in this activation process; this module is expressed by some *Sox2* -negative cells, but these cells express all the genes from the Wnt module (module 14). Similar to the DC module (module 15), module 16 co-localizes with module 1 from the E14.5 WT sample, while the localized pattern is absent in the E13.5 WT sample (Fig. 5B). RNA fluorescent in situ hybridization (FISH) validated this observation, showing that *Gal*, a marker featured in this module, is present in the upper dermis region in E13.5 SmoM2 but not in the paired WT, while *Gal* -positive cells are concentrated in the DC in E14.5 WT (Fig. S13A and S13C). We also noticed that the Wnt level (represented by *Lef1* expression levels) of this group of cells is comparable to that of DC cells, which are significantly higher than other cells involved in Wnt signaling (Fig. 5C, S13B and S13D). This suggests that module 16 may highlight cells committed to transitioning to DC states, potentially induced by high Wnt and SHH levels.

Using this example, we demonstrate the ability of LMD to capture gene modules that depict different cell states within a biological process that is represented by incremental activation of genes.

#### LMD detects a novel mutant-specific cell state

We identified a gene module from E13.5 SmoM2, module 12, showing a localized pattern in both E13.5 and E14.5 SmoM2, which is absent in their paired WT samples (Fig. 6A). The functional connection between two localized patterns is further validated by the co-localization of module 12 with module 15 from E14.5 SmoM2 when projecting module 12 onto E14.5 SmoM2, with a Jaccard index of 0.872. They also share 3 of the top 5 significantly enriched pathways from MsigDB, including p53 pathway (Fig. S12). Additionally, we validate this finding using FISH, demonstrating that *Trp53inp1*, a marker in both E13.5 SmoM2 module 12 and E14.5 SmoM2 module 15, is present in the upper dermis region in E13.5 and E14.5 SmoM2 but expressed at a negligible level in the paired WT (Fig. S13A and S13C).

We observed that module 12 delineates two distinct cell populations in E13.5 SmoM2: one predominantly in the G1 phase and the other in the G2M phase, with few cells in the S phase. When we regressed out the cell cycle score (see Section 4.3), these two populations merged, indicating that their separation was primarily driven by cell cycle differences (Fig. 6A, first row). A similar result is observed in E14.5 SmoM2, where module 12 also consists of two distinct cell populations (one in G1 and the other in G2M) that merge after regressing out the cell cycle score (Fig. 6A, second row). Given the phase composition and the significant enrichment of the p53 pathway in module 12 (Fig. S12), we hypothesize that module 12 delineates a quiescent cell state with G1 cells being the immediate progeny of a cell division (G2M). We test this hypothesis using FISH and EdU pulse-chase analysis. We show that most E14.5 *Trp53inp1* -positive cells do not incorporate EdU nucleotide, indicating they are not in the S phase of the cell cycle and largely quiescent. However, when pulsed with EdU 24 hours prior, we found that many precursors are labeled with EdU, suggesting that these cells are the progeny of proliferating cells (Fig. 6B). Additionally, the EdU percentage of *Trp53inp1* -positive cells is significantly higher when pulsed with EdU 24 hours prior either at E12.5 (chase to E13.5) and E13.5 (chase to E14.5) (Fig. 6C), indicating that most *Trp53inp1* -positive cells are quiescent but represent immediate progeny of a cell division. Thus, module 12 represents a mutant-specific quiescent cell state that represents the transition from proliferation to quiescence, a process that is normally coupled to DC differentiation. This transition from proliferation to quiescence occurs continuously between embryonic day 13.5 and 14.5. Since this module is specific to SmoM2, it may suggest a regulatory mechanism of cell cycle exit influenced by SHH pathway.

In this instance, we show that LMD can capture gene modules delineating condition-specific cell subsets without needing sample integration or differential abundance testing, effectively avoiding the need to use “batch correction” methods that cannot truly separate batch effects from biological effects and thus are not reliable.

## 3 Discussion

We present LMD, a novel approach for identifying localized markers at multiple length scales in single-cell RNA sequencing data. The core idea of LMD is that the dynamics of a diffusion process for each gene on a given cell-cell affinity graph reflects certain properties of the gene expression pattern such as locality. We have shown that expression patterns concentrated over a compact region of the cell-cell affinity graph yield a slower progression of diffusion at the beginning of this process. This property can be represented by LMD-score for each gene, and it enables the prioritization of localized expression patterns.

LMD consistently recovered known cell-type markers from several mouse and human scRNA-seq data, often outperforming other marker identification algorithms, which do not necessarily prioritize the localization of candidate markers. This success implies that many known markers are inherently localized, highlighting the necessity for algorithms like LMD that can distinguish between local and global expression profiles. Additionally, LMD’s top markers also enhance the separation of cells into distinct clumps. We further demonstrate the advantages of markers exhibiting localized patterns through two biological examples: LMD’s markers can identify compact groups of similar cells regardless of scale. Additionally, LMD’s markers can be used to match functionally similar cell populations between samples and detect condition-specific cell subsets without requiring sample integration, effectively avoiding the need to use “batch correction” methods that cannot truly separate batch effects from biological effects and thus are not reliable. LMD achieves these results without relying on existing cell annotations or gene functions. Its candidate markers are selected solely based on their localization within the cell-cell affinity graph.

In our second example, the mouse hair follicle dermal condensates system, we identified a previously uncharacterized potential intermediate state committed to DC, with *Gal* as a featured marker for this cell state. We also discovered a novel mutant-specific cell population located in two distinct regions within a single cluster in the UMAP visualization, with *Trp53inp1* as a marker. We demonstrated that this separation is driven by cell cycle differences and confirmed the biological connection through FISH and EdU pulse-chase experiments. These results suggest that this population may represent a quiescent cell state, potentially regulated by the SHH pathway. This example highlights the advantage of identifying markers in a cluster-independent manner, as traditional methods based on clustering and differential expression are less effective in single-cluster cases and in capturing connections between different cell groups.

However, a limitation of LMD is its potential bias introduced by the cell graph. Genes expressed in well-connected neighborhoods diffuse faster than those in more isolated neighborhoods of the same size. Further improvement of LMD will focus on better balancing gene distribution on the cell graph with the graph’s geometry. Additionally, using gene expression data for both constructing the cell graph and ranking the genes may introduce a bias toward highly variable genes. This bias arises because the PC coordinates used in graph construction tend to preserve information depicting the large variability of the data.

In this study, we demonstrated the effectiveness of LMD with scRNA-seq data. We expect that LMD can be adapted to other single-cell techniques, such as scATAC-seq and spatial transcriptomics.

## 4 Methods

### 4.1 Localized Marker Detector (LMD)

#### Input data

The input to LMD is a gene by cell log-normalized count matrix *X* ∈ ℝ^*m*×*n*^, where *G* is the set of all genes (|*G*| = *m*), *C* is the set of all cells (|*C*| = *n*), and the element *x*_*i,j*_ is the measured expression level of gene *i* in cell *j*.

#### Cell-cell affinity matrix *P*

First, we identify the *k*-nearest neighbors for each cell from the Euclidean distance between the transcription profiles after applying dimensional reduction, e.g. PCA (see Section 4.3). Next, the adjacency matrix *A* is constructed such that the element *a*_*i,j*_ is 1 if cell *i* is a *k*-nearest neighbor of cell *j* and 0 otherwise. Finally, the doubly-stochastic cell-cell affinity matrix *P* is obtained by applying the Sinkhorn-Knopp algorithm [49] to the symmetrized matrix (*A* + *A*^*T*^)/2.

#### Diffusion process

Given an initial state over the cells, propagate it over the nodes of the graph using a diffusion process. The initial state for gene *i* on the cell-cell affinity graph is the (*n* × 1) vector *ρ*^(*i*,0)^ whose elements are 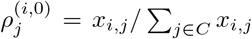. The state of gene *i* at time *t* is estimated using a diffusion process: *ρ*^(*i,t*)^ = (*P*^*t*^)^*′*^*ρ*^(*i*,0)^. Note that for *t* → ∞, *ρ*^(*i,t*)^ → 1/*n* · **1**, ∀*i* ∈ *G*.

#### LMD score

For a given gene *i*, the Kullback-Leibler divergence (KL-divergence) between the initial state and the state at time *t* is 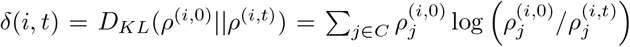. The Normalized KL-divergence is computed as *d*(*i, t*) = δ(*i, t*)/δ(*i*, ∞). We define the LMD score as LMDS = Σ_*t*_ *d*(*i, t*). In LMD, we compute LMDS over *t* = 2^*τ*^, *τ* = 1 … *T* for a *T* where the diagonal entries of 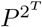 converge to 1/*n* with a maximum relative error of 1%. We rank candidate genes by ascending values of LMDS.

### 4.2 Gene modules

#### Identification

To identify gene modules, we selected the top candidate genes based on the knee point in the gene LMD score curve. The knee point is identified as the data point with the maximum perpendicular distance from the line connecting the first and last points[50]. For these selected genes, we computed the gene-gene pairwise distance matrix using the Jaccard distance [51] between denoised gene expression profiles across all cells, with denoising performed by ALRA [52]. A small Jaccard distance suggests that these two genes are co-localized in the cell graph. We then adopted a widely accepted strategy for gene module identification [53] where we processed the gene-gene pairwise distance matrix with hierarchical clustering with average linkage (`hclust` function from `stats` R package (v. 4.1.3) [54]) and we determined gene modules using the dynamic tree-cutting algorithm (`cutreeDynamic` function from `dynamicTreeCut` R package (v.1.63.1) [55]). We then removed outlier genes following this spectral filtering procedure[56]:

1. compute the gene-gene Jaccard Index matrix;
2. compute the first eigenvector and the associated eigenvalue *λ*_1_ of the Jaccard Index matrix;
3. compute the absolute values of the entries in the first eigenvector of the Jaccard Index matrix;
4. scale the absolute values by 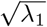
5. discard genes corresponding to scaled entries smaller than 0.5.

Only modules containing at least 10 genes were retained for downstream analysis.

#### Module Activity Scores

The module activity score (or module score) represents the probability of a module being expressed in a given cell, ranging from 0 to 1. To estimate this score, we performed the following sampling procedure 100 times to ensure stability: 1. Randomly select half of the genes from the module without replacement. 2. Calculate the average expression of these genes in each cell and cluster the cells using `GMM` function from `ClusterR` R package (v. 1.3.1)[57], with the optimal number of clusters estimated by `Optimal_Clusters_GMM` from `ClusterR` using the `BIC` criterion. 3. Assign the resulting clusters into two categories (0 - not expressing the module; 1 - expressing the module) by performing hierarchical clustering on the cluster centroids with average linkage (`hclust` function from `stats` R package), then cutting the tree into two clusters (`cutree` function from `stats` R package). Finally, determine the module activity score for each cell as the proportion of times it was assigned to category 1 over 100 iterations.

#### Cell type-specific activated modules

Cell type-specific activated modules are those for which the Jaccard index between the module score and the one-hot vector representation of a given cell type exceeds 0.4.

#### Measuring co-localization between two gene modules

The degree of co-localization between two gene modules is quantified using the Jaccard index, which compares their module score vectors. A higher Jaccard index indicates greater co-localization between the two modules.

### 4.3 Experiments and analyses

#### Dataset and Processing

##### Tabula Muris

We extracted publicly available scRNA-seq mouse bone marrow (FACS and droplet-based), lung (FACS-based), and pancreas (FACS-based) datasets (see Section 4.4). For LMD input, we constructed a cell-cell kNN (*k* = 5) graph using the top 20 PCs coordinates provided in the original datasets. Genes expressed in more than 10 cells but fewer than 50% of cells were included for analysis. The raw count matrix was log-normalized using the `Seurat v4` R package[3]. For gene module identification in the Tabula Muris bone marrow FACS dataset, we used the `cutreeDynamic` function with the parameters `minClusterSize = 10` and `deepSplit = 1`. For visualization, we use t-SNE coordinates provided in the original datasets. For benchmark methods (see Section 4.3), we used the same data, scaled or transformed as required by each method’s input specifics, e.g. either raw count matrix, log-normalized count matrix, or PC coordinates.

##### Azimuth demo dataset

We extracted publicly available scRNA-seq datasets provided by Azimuth[3] and annotated them using the Azimuth app at a reasonable resolution level (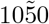 cell types). The datasets include: human motor cortex (annotated by subclass), mouse motor cortex (subclass), human pancreas (annotation.l1), human kidney (annotation.l2), human bone marrow (celltype.l2), and human lung v2 (ann level 3) (see Section 4.4). We removed cell types containing fewer than 10 cells. Following standard normalization, selection of the top 2, 000 highly variable genes, and scaling with the `Seurat v4` R package, we performed PCA and retained the top 20 PCs. For LMD input, we constructed a cell-cell kNN (*k* = 5) graph using the top 20 PCs coordinates. Genes expressed in more than 10 cells but fewer than 50% of cells were included for analysis. The raw count matrix was log-normalized using the `Seurat v4` R package. For benchmark methods, we used the same data, scaled or transformed as required by each method’s input specifics, e.g. either raw count matrix, log-normalized count data, or PC coordinates.

##### Mouse embryo skin data

We separated dermal cell populations from newly collected mouse embryo skin samples (see Section 4.3; aligned to the mouse genome mm10 using CellRanger (v.5.0.1)). To avoid batch effects from pooling or integrating, we analyzed each condition separately: E13.5 SmoM2, E13.5 WT, E14.5 SmoM2, and E14.5 WT. For each condition, we performed standard normalization, selected the top 2,000 highly variable genes, and scaled the data using the `Seurat v4` R package. We then applied PCA, retaining the number of PCs determined by the elbow plot[58]: E13.5 SmoM2 (14 PCs), E13.5 WT (12 PCs), E14.5 SmoM2 (12 PCs), and E14.5 WT (11 PCs). For LMD input in each condition, we constructed a cell-cell kNN (k=5) graph using the retained PC coordinates. Genes expressed in more than 10 cells but fewer than 50% of cells were included for analysis. The count data was log-normalized using the `Seurat v4` R package. For gene module identification, we used `cutreeDynamic` with `minClusterSize=10` and `deepSplit=2` for E13.5 SmoM2, and `deepSplit=1` for E13.5 WT, E14.5 SmoM2, and E14.5 WT. For visualization, we used UMAP coordinates generated from the retained PC coordinates.

#### Gold-standard cell-type marker sets

We employ two distinct gold-standard marker sets, as suggested in [12]. The first set includes marker genes listed in marker databases: for the Tabula Muris dataset, we use the CellMarker database[59] specific to the relevant tissue; for the Azimuth dataset, we use the marker list[19] used to annotate the reference dataset at the annotation level described in Section 4.3. The second set consists of the top 100 genes with the maximum differential expression across cell types, using cell-type annotations provided by the original studies (Tabula Muris) or annotated by Azimuth (Azimuth dataset).

#### Gene Ranking by Competing Method

LMD was qualitatively compared to other marker gene identification methods: singleCellHaystack, Hotspot, SEMITONES, MarcoPolo, Seurat v4, and HighlyVariableGenes. Each method uses the same gene list as input. The gene rankings for each method are summarized below:

– **singleCellHaystack**: We use singleCellHaystack1.0[60] with the advanced mode of the highD method. Genes are ranked in ascending order of log(adjusted p-value).
– **Hotspot**: Genes are ranked in ascending order of FDR.
– **SEMITONES**: Genes are ranked in ascending order of their minimum adjusted p-value in any reference cell. If two genes had the same adjusted p-value, we prioritize genes with larger absolute enrichment scores in any reference cell.
– **MarcoPolo**: The gene rankings is given by MarcoPolo rank.
– **Seurat v4**: The cluster resolution is set to 1.5. We used the `wilcoxauc.Seurat` function from `presto` R package (v.1.0.0)[61] to perform the fast Wilcoxon rank sum test among clusters. Genes are ranked in ascending order based on their minimum adjusted p-value among clusters. In cases where two genes had the same adjusted p-value, genes with larger fold-change were prioritized.
– **HighlyVariableGenes**: Genes are ranked by `FindVariableFeatures` function with the `VST` option implemented in Seurat v4.

#### Robustness test

LMD is based on the cell-cell kNN graph, where we set the default input space as top 20 PCs and the default value of *k* = 5. Using the Tabula Muris bone marrow FACS dataset[62], we show that the choice of *k* and input spaces has no significant effect on our results. First, we test the effect of different *k*(*k* = 3, 4, 5, 6, 7, 8, 9, 10, 20, 30, 40, 50) while fixing the input space (top 20 PCs). We assess the rank stability by computing the Jaccard index between the top *N* genes using the default *k* = 5 and different *k* (Fig. S6A). We also test the stability of AUROC across different *k* when evaluating LMD’s performance on the two gold-standard gene sets described in Section 4.3 (Fig. S6B). Next, we test the effect of different input spaces (top 10, 20, 30, 40, 50, 75, 100 PCs, t-SNE) while fixing *k* = 5. We assess the rank stability by comparing the top *N* genes using the default top 20 PCs and various input spaces (Fig. S6C). Additionally, we test the stability of AUROC across different input spaces on the same gold-standard gene sets (Fig. S6D).

#### Scalability

We assessed the scalability of LMD by running it on nine datasets (Fig. 3), with cell numbers ranging from 1,500 to 30,000 and gene numbers from 13,000 to 34,000 on a Linux operating system with 72 2.70GHz cores. (Table S2). The runtime was primarily dependent on the number of cells rather than the number of genes. For a fixed cell number, runtimes were similar whether gene counts were around 14,000 or 34,000. The runtime increased with the number of cells: approximately 30 seconds for 1,500 cells and 44 minutes for 30,000 cells.

#### GO and Reactome Enrichment analysis

We perform Gene Ontology (GO) enrichment analysis using the `clusterProfiler` R package (v.4.2.2)[63], Reactome Pathway Database enrichment analysis with the `ReactomePA` R package (v.1.38.0)[64] and Molecular Signatures Database (MsigDB) enrichment analysis with `msigdbr` R package (v.7.5.1)[65] and `clusterProfiler` R package (v.4.2.2). The background gene set comprises all genes that are applied during the localized marker ranking process. We regard GO terms and pathways with Benjamini-Hochberg adjusted p-values smaller than 0.05 as significantly enriched.

#### Cell cycle annotation, embedding, and regression

We first extract the 2019-updated human cell cycle gene set[66] from `cc.genes.updated.2019` in `Seurat` v4 R package[3] and transform it to mouse genes using the `gprofiler2` R package (v.0.2.3)[67]. This reference cell cycle gene list, containing 54 G2/M phase markers and 41 S phase markers, is then used for annotation and embedding. Then we perform cell cycle scoring and annotation using the `Seurat` v4 R package. The `CellCycleScoring` function assigns cell cycle scores based on the reference cell cycle gene list, classifying cells into G1, S, or G2/M phases. To create UMAP/t-SNE embeddings that preserve only the heterogeneity from the cell cycle, we first run PCA on the reference cell cycle gene list, retaining the number of PCs determined by the elbow plot[58]. We then apply UMAP/t-SNE on the retained PCs. To create UMAP/t-SNE embeddings that subtract the heterogeneity from the cell cycle, we follow the `Seurat` workflow. We regress out cell cycle scores during data scaling, then perform dimensional reduction on the scaled data.

#### Experimental details of mouse embryo skin sample preparation

##### Mice

Axin2CreER[68] mice were bred to Rosa-lox-STOP-lox-SmoM2YFP[69] mice. A random population of both male and female embryos was used for all experiments. All procedures involving animal subjects were performed under the approval of the Institutional Animal Care and Use Committee of the Yale School of Medicine.

##### EdU and Tamoxifen administration

To induce expression of SmoM2YFP, tamoxifen (60*µg/gm* weight; Sigma Aldrich Cat#T5648) was administered to pregnant mice at E11.5 by oral gavage. To assess proliferation, EdU (5-ethynyl-2-deoxyuridine; Thermo Fisher Scientific Cat#E10187) was administered to pregnant mice intraperitoneally (25*µg/gm*) and embryos were collected after either 1.5*h* or 24 hours.

##### In situ hybridization

Formalin-fixed paraffin-embedded (FFPE) whole embryos were used for histological analysis. FFPE specimens were sectioned at 10*µm* thickness. The RNAscope Multiplex Fluorescent Detection Kit v2 (ACDBio, 323110) was used for single-molecule fluorescence in situ hybridization (FISH) according to the manufacturer’s protocol. Briefly, sections were deparaffinized and permeabilized with hydrogen peroxide followed by antigen retrieval and protease treatment before probe hybridization. After hybridization, amplification and probe detection were done using the Amp 1–3 reagents. Probe channels were targeted using the provided HRP-C1-3 reagents and tyramide signal amplification VIVID fluorophores—650 and 570 (ACD biotechne, 323271). EdU staining was done using the Click-it EdU Imaging Kit Alexa 488 (Life Technologies, c10338) according to the manufacturer’s instructions. Nuclear counter-stain was done using Hoechst 33342 (Invitrogen, H3570) before mounting with SlowFade Mountant. RNA scope probes used (ACDBio)—Mm-Lef1 (441861), Mm-Gal (400961) and Mm-Trp53inp1 (1161531).

##### Microscopy

FISH paraffin-embedded images were acquired using the Leica Stellaris 8 DMi8 confocal microscope with a ×40 oil immersion (Numerical Aperture 1.3) objective lens, scanned at 5 *µm* thickness, 1, 024 × 1, 024 pixel width, 400*Hz*.

##### Single-cell dissociation

Dorsolateral/flank skin was microdissected from E13.5 or E14.5 littermate control and mutant embryos and dissociated into a single-cell suspension using 0.25% trypsin (Gibco, Life Technologies) for 20 min at 37 °C. After genotyping, two to three embryos were pooled by condition. Single-cell suspensions were then stained with DAPI (Thermo Fisher Scientific, NBP2-31156) before fluorescence-activated cell sorting.

##### Fluorescence-activated cell sorting

DAPI-excluded live skin cells were sorted on a BD FACS Aria II (BD Biosciences) sorter with a 100 *µm* nozzle. Cells were sorted in bulk and submitted for 10X Genomics library preparation at 0.75 *−*1.0 × 10^6^*ml*^−1^ concentration in 4% fetal calf serum/phosphate-buffered saline (FCS/PBS) solution.

##### H-score quantification

For quantification based on FISH, cells with 4–5 dots were considered positive (according to the RNAScope manufacturer’s instructions) and subsections from a total of *n* = 4 different embryos were examined. To measure RNA expression levels, H scores were calculated according to ACDBio manufacturer’s instructions: a cell with 0 dot is scored 0, 1–3 dots is scored 1, 4–9 dots is scored 2, 10–15 dots and/or less than 10% clustered dots is scored 3 and more than 15 dots and/or more than 10% clustered dots is scored 4; then the final H score of a given cell type A is calculated by summing the (% cells scored B within all cells in A) × B for score B in 0–4. Bin 4 *Trp53inp1* + cells were quantified for %EdU.

##### scRNA-seq and library preparation

Chromium Single Cell 3’ GEM Library and Gel Bead Kit v3.1 (PN1000121) were used according to the manufacturer’s instructions in the Chromium Single Cell 3’ Reagents Kits V3.1 User Guide. After cDNA libraries were created, they were subjected to Novaseq 6000 (Illumina) sequencing. For each scRNA-seq experiment, control and littermate mutant samples were prepared in parallel at the same time, pooled and sequenced on the same lane.

### 4.4 Data availability

The Tabula Muris scRNA-seq datasets are available at [70]. These include FACS-based full-length transcript datasets for bone marrow [62], pancreas [71] and lung [72], as well as a microfluidic droplet-based 3’-end counting dataset for bone marrow [73]. The Azimuth demo datasets are available at [19]. These include human motor cortex[74], mouse motor cortex[75], human pancreas[76], human kidney[77], human bone marrow[78], and human lung v2[79]. The novel mouse embryonic skin dataset generated and analyzed in this study will be deposited in the Gene Expression Omnibus (GEO) before publication and the processed Seurat data objects are available from the corresponding authors upon reasonable request.

### 4.5 Code availability

The R package of LMD and the code used for data analysis will be available soon on GitHub.

## 4.6 Acknowledgment

The authors thank Ronald Coifman, Junchen Yang, Will Yaochen Zhu for fruitful discussions. Y.K. is supported by [R01GM131642, UM1PA051410, U54AG076043, U54AG079759, U01DA053628, P50CA121974, and R33DA047037]. X.C. is partially supported by NSF [DMS-2237842, DMS-2007040] and Simons Foundation. P.M. is supported by NIH/NIAMS [R01AR076420].

## References

1. J. S. Fleck, J. G. Camp, and B. Treutlein, “What is a cell type?,” Science, vol. 381, no. 6659, pp. 733–734, 2023.

2. C. Cheng, W. Chen, H. Jin, and X. Chen, “A review of single-cell rna-seq annotation, integration, and cell–cell communication,” Cells, vol. 12, no. 15, p. 1970, 2023.

3. Y. Hao, S. Hao, E. Andersen-Nissen, W. M. Mauck, S. Zheng, A. Butler, M. J. Lee, A. J. Wilk, C. Darby, M. Zager, et al., “Integrated analysis of multimodal single-cell data,” Cell, vol. 184, no. 13, pp. 3573–3587, 2021.

4. R. Patrick, D. T. Humphreys, V. Janbandhu, A. Oshlack, J. W. Ho, R. P. Harvey, and K. K. Lo, “Sierra: discovery of differential transcript usage from polya-captured single-cell rna-seq data,” Genome biology, vol. 21, no. 1, pp. 1–27, 2020.

5. V. Svensson, S. A. Teichmann, and O. Stegle, “Spatialde: identification of spatially variable genes,” Nature methods, vol. 15, no. 5, pp. 343–346, 2018.

6. Z. Miao, K. Deng, X. Wang, and X. Zhang, “Desingle for detecting three types of differential expression in single-cell rna-seq data,” Bioinformatics, vol. 34, no. 18, pp. 3223–3224, 2018.

7. S. Nabavi, D. Schmolze, M. Maitituoheti, S. Malladi, and A. H. Beck, “Emdomics: a robust and powerful method for the identification of genes differentially expressed between heterogeneous classes,” Bioinformatics, vol. 32, no. 4, pp. 533–541, 2016.

8. K. D. Korthauer, L.-F. Chu, M. A. Newton, Y. Li, J. Thomson, R. Stewart, and C. Kendziorski, “A statistical approach for identifying differential distributions in single-cell rna-seq experiments,” Genome biology, vol. 17, pp. 1–15, 2016.

9. M. D. Robinson, D. J. McCarthy, and G. K. Smyth, “edger: a bioconductor package for differential expression analysis of digital gene expression data,” bioinformatics, vol. 26, no. 1, pp. 139–140, 2010.

10. X. Qiu, Q. Mao, Y. Tang, L. Wang, R. Chawla, H. A. Pliner, and C. Trapnell, “Reversed graph embedding resolves complex single-cell trajectories,” Nature methods, vol. 14, no. 10, pp. 979–982, 2017.

11. A. Vandenbon and D. Diez, “A clustering-independent method for finding differentially expressed genes in single-cell transcriptome data,” Nature communications, vol. 11, no. 1, p. 4318, 2020.

12. C. Kim, H. Lee, J. Jeong, K. Jung, and B. Han, “Marcopolo: a method to discover differentially expressed genes in single-cell rna-seq data without depending on prior clustering,” Nucleic acids research, vol. 50, no. 12, pp. e71–e71, 2022.

13. A. H. C. Vlot, S. Maghsudi, and U. Ohler, “Cluster-independent marker feature identification from single-cell omics data using semitones,” Nucleic Acids Research, vol. 50, no. 18, pp. e107–e107, 2022.

14. D. DeTomaso and N. Yosef, “Hotspot identifies informative gene modules across modalities of single-cell genomics,” Cell systems, vol. 12, no. 5, pp. 446–456, 2021.

15. T. M. Consortium, O. coordination Schaum Nicholas 1 Karkanias Jim 2 Neff Norma F. 2 May Andrew P. 2 Quake Stephen R. quake@ stanford. edu 2 3 f Wyss-Coray Tony twc@ stanford. edu 4 5 6 g Darmanis Spyros spyros. darmanis@ czbiohub. org 2 h, L. coordination Batson Joshua 2 Botvinnik Olga 2 Chen Michelle B. 3 Chen Steven 2 Green Foad 2 Jones Robert C. 3 Maynard Ashley 2 Penland Lolita 2 Pisco Angela Oliveira 2 Sit Rene V. 2 Stanley Geoffrey M. 3 Webber James T. 2 Zanini Fabio 3, and C. data analysis Batson Joshua 2 Botvinnik Olga 2 Castro Paola 2 Croote Derek 3 Darmanis Spyros 2 DeRisi Joseph L. 2 27 Karkanias Jim 2 Pisco Angela Oliveira 2 Stanley Geoffrey M. 3 Webber James T. 2 Zanini Fabio 3, “Single-cell transcriptomics of 20 mouse organs creates a tabula muris,” Nature, vol. 562, no. 7727, pp. 367–372, 2018.

16. R. R. Coifman, S. Lafon, A. B. Lee, M. Maggioni, B. Nadler, F. Warner, and S. W. Zucker, “Geometric diffusions as a tool for harmonic analysis and structure definition of data: Multiscale methods,” Proceedings of the National Academy of Sciences, vol. 102, no. 21, pp. 7432–7437, 2005.

17. R. R. Coifman and M. Maggioni, “Diffusion wavelets,” Applied and computational harmonic analysis, vol. 21, no. 1, pp. 53–94, 2006.

18. C. J. Dsilva, R. Talmon, N. Rabin, R. R. Coifman, and I. G. Kevrekidis, “Nonlinear intrinsic variables and state reconstruction in multiscale simulations,” The Journal of chemical physics, vol. 139, no. 18, 2013.

19. Azimuth Demo Datasets, “Azimuth Demo Datasets.” https://azimuth.hubmapconsortium.org/references/, 2021.

20. B. Ranjan, W. Sun, J. Park, K. Mishra, F. Schmidt, R. Xie, F. Alipour, V. Singhal, I. Joanito, M. A. Honardoost, et al., “Dubstepr is a scalable correlation-based feature selection method for accurately clustering single-cell data,” Nature Communications, vol. 12, no. 1, p. 5849, 2021.

21. M. Gillespie, B. Jassal, R. Stephan, M. Milacic, K. Rothfels, A. Senff-Ribeiro, J. Griss, C. Sevilla, L. Matthews, C. Gong, et al., “The reactome pathway knowledgebase 2022,” Nucleic acids research, vol. 50, no. D1, pp. D687–D692, 2022.

22. B. Tandon, L. Peterson, J. Gao, B. Nelson, S. Ma, S. Rosen, and Y.-H. Chen, “Nuclear overexpression of lymphoid-enhancer-binding factor 1 identifies chronic lymphocytic leukemia/small lymphocytic lymphoma in small b-cell lymphomas,” Modern Pathology, vol. 24, no. 11, pp. 1433–1443, 2011.

23. L.-S. Lu, J. Tung, N. Baumgarth, O. Herman, M. Gleimer, L. A. Herzenberg, and L. A. Herzenberg, “Identification of a germ-line pro-b cell subset that distinguishes the fetal/neonatal from the adult b cell development pathway,” Proceedings of the National Academy of Sciences, vol. 99, no. 5, pp. 3007–3012, 2002.

24. Y.-S. Li, K. Hayakawa, and R. R. Hardy, “The regulated expression of b lineage associated genes during b cell differentiation in bone marrow and fetal liver.,” The Journal of experimental medicine, vol. 178, no. 3, pp. 951–960, 1993.

25. J. J. Peschon, P. J. Morrissey, K. H. Grabstein, F. J. Ramsdell, E. Maraskovsky, B. C. Gliniak, L. S. Park, S. F. Ziegler, D. E. Williams, C. B. Ware, et al., “Early lymphocyte expansion is severely impaired in interleukin 7 receptor-deficient mice.,” The Journal of experimental medicine, vol. 180, no. 5, pp. 1955–1960, 1994.

26. M. Jensen, G. Tan, S. Forman, A. M. Wu, and A. Raubitschek, “Cd20 is a molecular target for scfvfc: zeta receptor redirected t cells: implications for cellular immunotherapy of cd20+ malignancy,” Biology of blood and marrow transplantation, vol. 4, no. 2, pp. 75–83, 1998.

27. T. F. Tedder, J. C. Poe, and K. M. Haas, “Cd22: a multifunctional receptor that regulates b lymphocyte survival and signal transduction,” Advances in immunology, vol. 88, pp. 1–50, 2005.

28. J.-Y. Bonnefoy, S. Lecoanet-Henchoz, J.-F. Gauchat, P. Graber, J.-P. Aubry, P. Jeannin, and C. Plater-Zyberk, “Structure and functions of cd23,” International reviews of immunology, vol. 16, no. 1-2, pp. 113–128, 1997.

29. J. Sáez de Guinoa, L. Barrio, M. Mellado, and Y. R. Carrasco, “Cxcl13/cxcr5 signaling enhances bcr-triggered b-cell activation by shaping cell dynamics,” Blood, The Journal of the American Society of Hematology, vol. 118, no. 6, pp. 1560–1569, 2011.

30. Z. Liu, Y. Gu, S. Chakarov, C. Bleriot, I. Kwok, X. Chen, A. Shin, W. Huang, R. J. Dress, C.-A. Dutertre, et al., “Fate mapping via ms4a3-expression history traces monocyte-derived cells,” Cell, vol. 178, no. 6, pp. 1509–1525, 2019.

31. X. Xie, Q. Shi, P. Wu, X. Zhang, H. Kambara, J. Su, H. Yu, S.-Y. Park, R. Guo, Q. Ren, et al., “Single-cell transcriptome profiling reveals neutrophil heterogeneity in homeostasis and infection,” Nature immunology, vol. 21, no. 9, pp. 1119–1133, 2020.

32. D. M. Calcagno, C. Zhang, A. Toomu, K. Huang, V. K. Ninh, S. Miyamoto, A. D. Aguirre, Z. Fu, J. Heller Brown, and K. R. King, “Siglecf (hi) marks late-stage neutrophils of the infarcted heart: a single-cell transcriptomic analysis of neutrophil diversification,” Journal of the American Heart Association, vol. 10, no. 4, p. e019019, 2021.

33. K. J. Eash, A. M. Greenbaum, P. K. Gopalan, D. C. Link, et al., “Cxcr2 and cxcr4 antagonistically regulate neutrophil trafficking from murine bone marrow,” The Journal of clinical investigation, vol. 120, no. 7, pp. 2423–2431, 2010.

34. K. J. Goodall, A. Nguyen, A. Matsumoto, J. R. McMullen, S. B. Eckle, P. Bertolino, L. C. Sullivan, and D. M. Andrews, “Multiple receptors converge on h2-q10 to regulate nk and γδt-cell development,” Immunology and Cell Biology, vol. 97, no. 3, pp. 326–339, 2019.

35. K. H. Kwack, N. A. Lamb, J. E. Bard, E. D. Kramer, L. Zhang, S. I. Abrams, and K. L. Kirkwood, “Discovering myeloid cell heterogeneity in mandibular bone–cell by cell analysis,” Frontiers in Physiology, vol. 12, p. 731549, 2021.

36. S. Ping, X. Qiu, M. E. Gonzalez-Toledo, X. Liu, and L.-R. Zhao, “Stem cell factor in combination with granulocyte colony-stimulating factor protects the brain from capillary thrombosis-induced ischemic neuron loss in a mouse model of cadasil,” Frontiers in Cell and Developmental Biology, vol. 8, p. 627733, 2021.

37. R. Qu, K. Gupta, D. Dong, Y. Jiang, B. Landa, C. Saez, G. Strickland, J. Levinsohn, P.-l. Weng, M. M. Taketo, et al., “Decomposing a deterministic path to mesenchymal niche formation by two intersecting morphogen gradients,” Developmental cell, vol. 57, no. 8, pp. 1053–1067, 2022.

38. K. Gupta, J. Levinsohn, G. Linderman, D. Chen, T. Y. Sun, D. Dong, M. M. Taketo, M. Bosenberg, Y. Kluger, K. Choate, et al., “Single-cell analysis reveals a hair follicle dermal niche molecular differentiation trajectory that begins prior to morphogenesis,” Developmental cell, vol. 48, no. 1, pp. 17–31, 2019.

39. P. Myung, T. Andl, and R. Atit, “The origins of skin diversity: lessons from dermal fibroblasts,” Development, vol. 149, no. 23, p. dev200298, 2022.

40. R. S. Wu and W. M. Bonner, “Separation of basal histone synthesis from s-phase histone synthesis in dividing cells,” Cell, vol. 27, no. 2, pp. 321–330, 1981.

41. H. Liao, R. Winkfein, G. Mack, J. Rattner, and T. Yen, “Cenp-f is a protein of the nuclear matrix that assembles onto kinetochores at late g2 and is rapidly degraded after mitosis.,” The Journal of cell biology, vol. 130, no. 3, pp. 507–518, 1995.

42. C. Zhu, J. Zhao, M. Bibikova, J. D. Leverson, E. Bossy-Wetzel, J.-B. Fan, R. T. Abraham, and W. Jiang, “Functional analysis of human microtubule-based motor proteins, the kinesins and dyneins, in mitosis/cytokinesis using rna interference,” Molecular biology of the cell, vol. 16, no. 7, pp. 3187–3199, 2005.

43. M. K. Diril, C. K. Ratnacaram, V. Padmakumar, T. Du, M. Wasser, V. Coppola, L. Tessarollo, and P. Kaldis, “Cyclin-dependent kinase 1 (cdk1) is essential for cell division and suppression of dna re-replication but not for liver regeneration,” Proceedings of the National Academy of Sciences, vol. 109, no. 10, pp. 3826–3831, 2012.

44. R. S. Hames and A. M. Fry, “Alternative splice variants of the human centrosome kinase nek2 exhibit distinct patterns of expression in mitosis,” Biochemical Journal, vol. 361, no. 1, pp. 77–85, 2002.

45. A. Liberzon, C. Birger, H. Thorvaldsdottir, M. Ghandi, J. P. Mesirov, and P. Tamayo, “The molecular signatures database hallmark gene set collection,” Cell systems, vol. 1, no. 6, pp. 417–425, 2015.

46. K. Hovanes, T. W. Li, J. E. Munguia, T. Truong, T. Milovanovic, J. Lawrence Marsh, R. F. Holcombe, and M. L. Waterman, “β-catenin–sensitive isoforms of lymphoid enhancer factor-1 are selectively expressed in colon cancer,” Nature genetics, vol. 28, no. 1, pp. 53–57, 2001.

47. M. Filali, N. Cheng, D. Abbott, V. Leontiev, and J. F. Engelhardt, “Wnt-3a/β-catenin signaling induces transcription from the lef-1 promoter* 210,” Journal of Biological Chemistry, vol. 277, no. 36, pp. 33398–33410, 2002.

48. R. R. Driskell, A. Giangreco, K. B. Jensen, K. W. Mulder, and F. M. Watt, “Sox2-positive dermal papilla cells specify hair follicle type in mammalian epidermis,” Development, 2009.

49. R. Sinkhorn and P. Knopp, “Concerning nonnegative matrices and doubly stochastic matrices,” Pacific Journal of Mathematics, vol. 21, no. 2, pp. 343–348, 1967.

50. V. Satopaa, J. Albrecht, D. Irwin, and B. Raghavan, “Finding a” kneedle” in a haystack: Detecting knee points in system behavior,” in 2011 31st international conference on distributed computing systems workshops, pp. 166–171, IEEE, 2011.

51. S.-H. Cha, “Comprehensive survey on distance/similarity measures between probability density functions,” City, vol. 1, no. 2, p. 1, 2007.

52. G. C. Linderman, J. Zhao, M. Roulis, P. Bielecki, R. A. Flavell, B. Nadler, and Y. Kluger, “Zero-preserving imputation of single-cell rna-seq data,” Nature communications, vol. 13, no. 1, p. 192, 2022.

53. P. Langfelder and S. Horvath, “Wgcna: an r package for weighted correlation network analysis,” BMC bioinformatics, vol. 9, no. 1, pp. 1–13, 2008.

54. R Core Team, R: A Language and Environment for Statistical Computing. R Foundation for Statistical Computing, Vienna, Austria, 2022.

55. P. Langfelder, B. Zhang, and with contributions from Steve Horvath, dynamicTreeCut: Methods for Detection of Clusters in Hierarchical Clustering Dendrograms, 2016. R package version 1. 63–1.

56. F. Parisi, F. Strino, B. Nadler, and Y. Kluger, “Ranking and combining multiple predictors without labeled data,” Proceedings of the National Academy of Sciences, vol. 111, no. 4, pp. 1253–1258, 2014.

57. L. Mouselimis, ClusterR: Gaussian Mixture Models, K-Means, Mini-Batch-Kmeans, K-Medoids and Affinity Propagation Clustering, 2023. R package version 1.3.1.

58. Harvard Chan Bioinformatics Core, “Elbow plot: quantitative approach.” https://hbctraining.github.io/scRNA-seq/lessons/elbow_plot_metric.html, 2021.

59. X. Zhang, Y. Lan, J. Xu, F. Quan, E. Zhao, C. Deng, T. Luo, L. Xu, G. Liao, M. Yan, et al., “Cellmarker: a manually curated resource of cell markers in human and mouse,” Nucleic acids research, vol. 47, no. D1, pp. D721–D728, 2019.

60. A. Vandenbon and D. Diez, “A universal tool for predicting differentially active features in single-cell and spatial genomics data,” Scientific Reports, vol. 13, no. 1, p. 11830, 2023.

61. I. Korsunsky, A. Nathan, N. Millard, and S. Raychaudhuri, presto: Fast Functions for Differential Expression using Wilcox and AUC, 2024. R package version 1.0.0.

62. Tabula Muris Consortium, “Bone marrow FACS dataset.” https://figshare.com/ndownloader/files/13092380, 2018.

63. T. Wu, E. Hu, S. Xu, M. Chen, P. Guo, Z. Dai, T. Feng, L. Zhou, W. Tang, L. Zhan, et al., “clusterprofiler 4.0: A universal enrichment tool for interpreting omics data,” The innovation, vol. 2, no. 3, 2021.

64. G. Yu and Q.-Y. He, “Reactomepa: an r/bioconductor package for reactome pathway analysis and visualization,” Molecular BioSystems, vol. 12, no. 2, pp. 477–479, 2016.

65. I. Dolgalev, msigdbr: MSigDB Gene Sets for Multiple Organisms in a Tidy Data Format, 2022. R package version 7.5.1.9001.

66. I. Tirosh, B. Izar, S. M. Prakadan, M. H. Wadsworth, D. Treacy, J. J. Trombetta, A. Rotem, C. Rodman, C. Lian, G. Murphy, et al., “Dissecting the multicellular ecosystem of metastatic melanoma by single-cell rna-seq,” Science, vol. 352, no. 6282, pp. 189–196, 2016.

67. L. Kolberg, U. Raudvere, I. Kuzmin, J. Vilo, and H. Peterson, “gprofiler2–an r package for gene list functional enrichment analysis and namespace conversion toolset g: Profiler,” F1000Research, vol. 9, 2020.

68. R. Van Amerongen, A. N. Bowman, and R. Nusse, “Developmental stage and time dictate the fate of wnt/β-catenin-responsive stem cells in the mammary gland,” Cell stem cell, vol. 11, no. 3, pp. 387–400, 2012.

69. J. Jeong, J. Mao, T. Tenzen, A. H. Kottmann, and A. P. McMahon, “Hedgehog signaling in the neural crest cells regulates the patterning and growth of facial primordia,” Genes & development, vol. 18, no. 8, pp. 937–951, 2004.

70. Tabula Muris Consortium, “Tabula Muris consortium dataset.” https://tabula-muris.ds.czbiohub.org/, 2018.

71. Tabula Muris Consortium, “Pancreas FACS dataset.” https://figshare.com/ndownloader/files/13092386, 2018.

72. Tabula Muris Consortium, “Lung FACS dataset.” https://figshare.com/ndownloader/files/13092194, 2018.

73. Tabula Muris Consortium, “Bone marrow droplet dataset.” https://figshare.com/ndownloader/files/13089821, 2018.

74. Azimuth Demo Datasets, “Human - Motor Cortex demo dataset.” https://seurat.nygenome.org/azimuth/demo_datasets/allen_m1c_2019_ssv4.rds, 2021.

75. Azimuth Demo Datasets, “Mouse - Motor Cortex demo dataset.” https://seurat.nygenome.org/azimuth/demo_datasets/allen_mop_2020.rds, 2021.

76. Azimuth Demo Datasets, “Human - Pancreas demo dataset.” https://seurat.nygenome.org/azimuth/demo_datasets/enge.rds, 2021.

77. Azimuth Demo Datasets, “Human - Kidney demo dataset.” https://seurat.nygenome.org/azimuth/demo_datasets/kidney_demo_stewart.rds, 2021.

78. Azimuth Demo Datasets, “Human - Bone Marrow demo dataset.” https://seurat.nygenome.org/azimuth/demo_datasets/bmcite_demo.rds, 2021.

79. Azimuth Demo Datasets, “Human - Lung v2 (HLCA) demo dataset.” https://seurat.nygenome.org/hlca_ref_files/ts_opt.rds, 2021.

