## Supplemental Figure 1-14; Table 1-2; Additional data for "LMD: Cluster-Independent Multiscale Marker Identification in Single-cell RNA-seq Data"

#### S1. Supplementary Figures

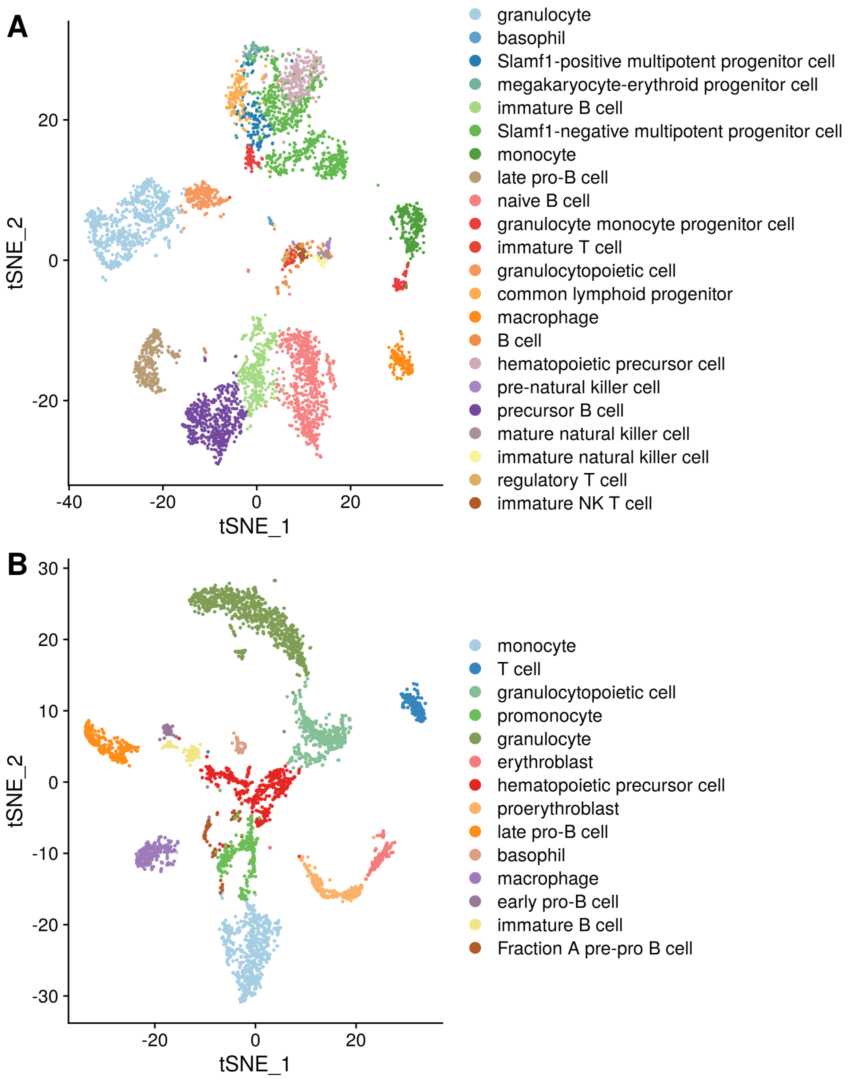


**Fig. S1.** t-SNE of two mouse bone marrow datasets, color-coded by pre-annotated cell types. (A) FACS-based: 5,037 cells across 22 cell types. (B) Droplet-based: 3,652 cells across 14 cell types.

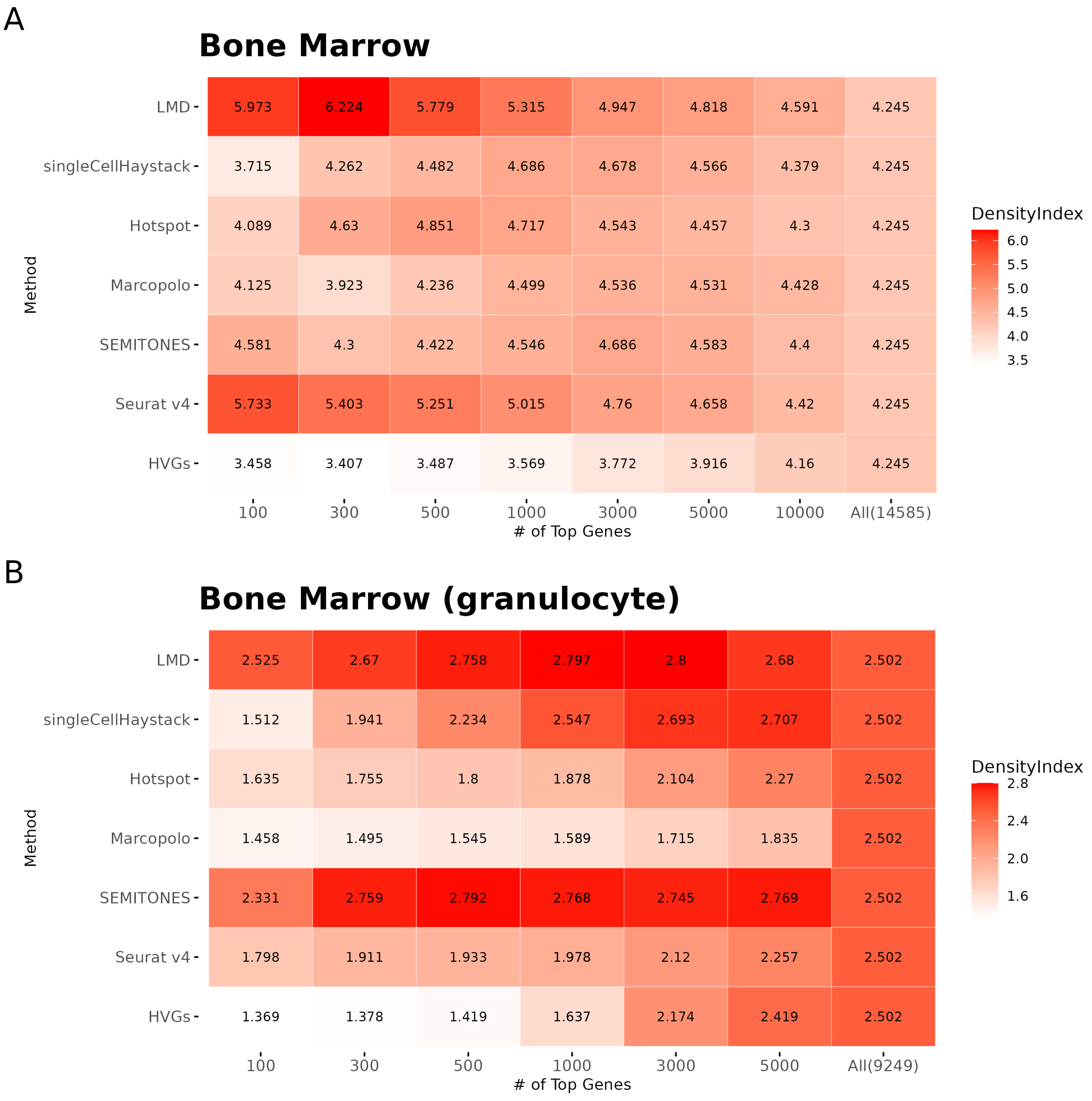


**Fig. S2.** Cluster Separability Comparison between LMD and other methods on (A) the mouse Bone Marrow dataset and (B) the granulocyte subset of the mouse Bone Marrow dataset. Separability is measured by the density index, computed over the top 20 PCs of cells expressing at least one gene from the top-ranked gene lists identified by LMD, singleCellHaystack, Hotspot, Marcopolo, SEMITONES, Seurat v4, and Highly Variable Genes.

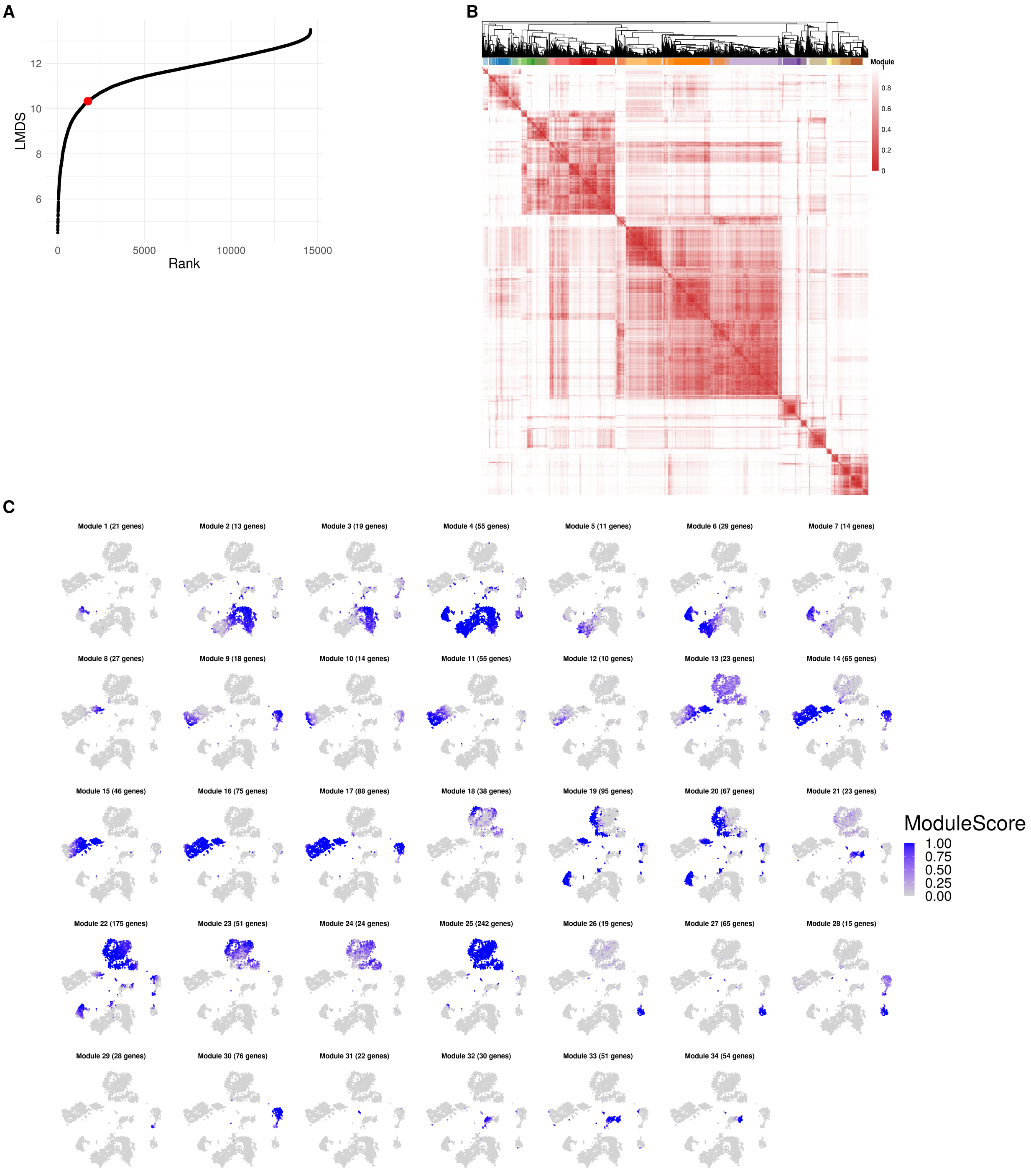


**Fig. S3.** Obtain gene modules of mouse bone marrow dataset. (A) Define a cutoff for the number of localized genes: The knee point (red dots) in the gene LMD scores curve indicates a cutoff at 1741 genes. (B) Gene-gene Jaccard distance among the selected localized genes, organized through hierarchical clustering. The color bar represents 34 distinct modules, obtained by dynamically cutting the dendrogram and filtering out outlier genes for each module. (C) t-SNE of 34 gene modules, color-coded by module score.

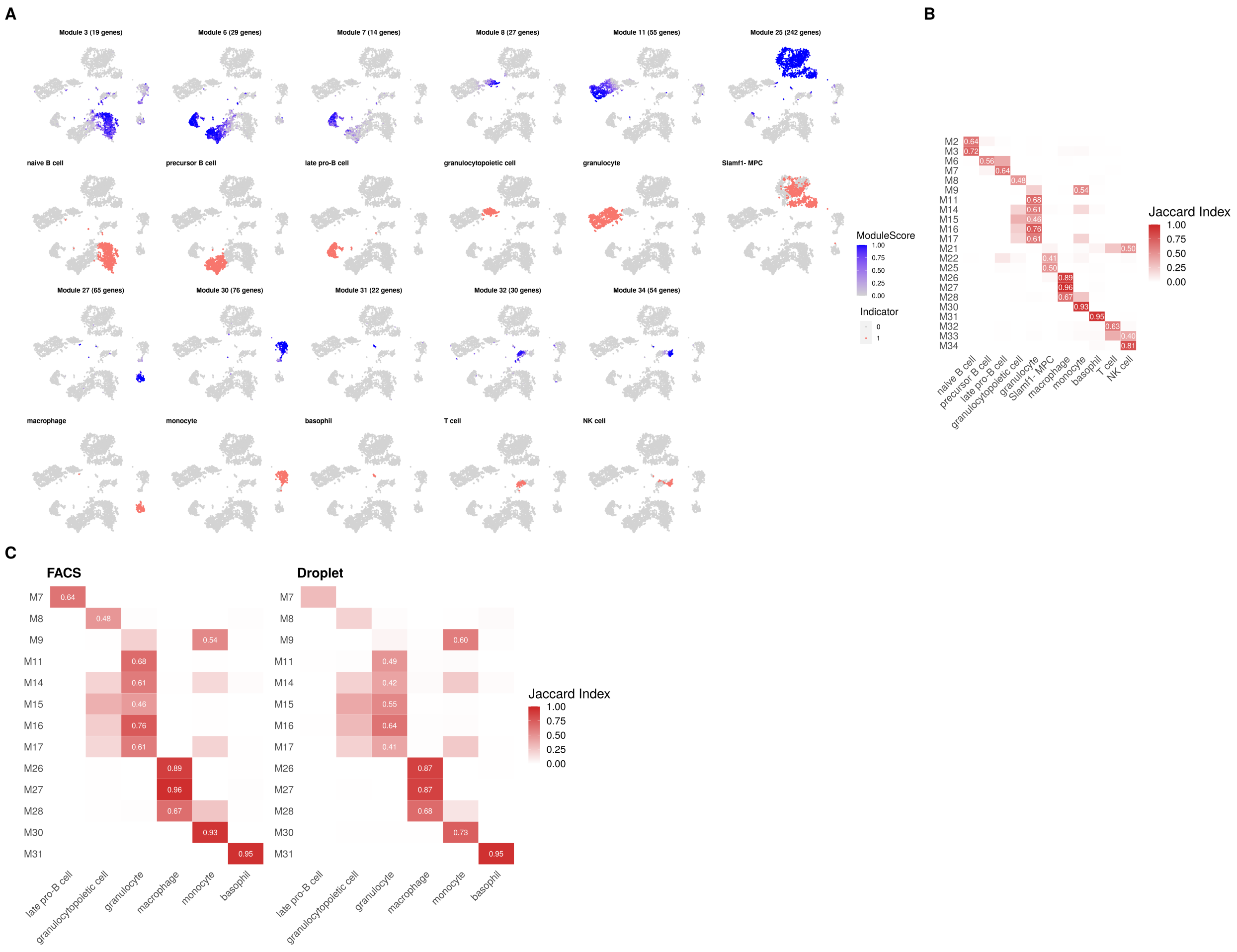

**Fig. S4.** Aligning gene modules with pre-annotated cell types. (A) Gene modules with the highest association to each cell type, determined by Jaccard index, color-coded by module score (first and third rows) or one-hot cell type indicator (second and fourth rows). The associations are as follows, with the genes used as canonical markers for annotation by the Tabula Muris group[1]. Module 3 (*Fcer2a*) - Naive B cell, module 6 (*Rag2*) - Precursor B cell, module 7 (*Lef1*) - Late pro-B cell, module 8 (*Ms4a3*) - Granulocytopoietic cell, module 11 (*Mmp9*) - Granulocyte, module 25 (*Cd34*, *Gpr56*, *Kit*) - Slamf1-MPC, module 27 (*Siglech*) - Macrophage, module 30 (*Emr1*) - Monocyte, module 31 (*Fcer1a*, *Mcpt8*) - Basophil, module 32 (*Cd3e*, *Cd8a*) - T cell, module 34 (*Klrb1a*, *Klrb1b*, *Klrb1c*, *Ncr1*) - NK cell. (B) Jaccard index of 22 cell type-specific activated modules (rows) across 11 cell types (columns). (C) Cross-dataset verification of module specificity: Jaccard index of 13 cell type-specific activated gene modules (rows) across 6 cell types (columns) in two bone marrow datasets FACS-based (left) and Droplet-based (right).

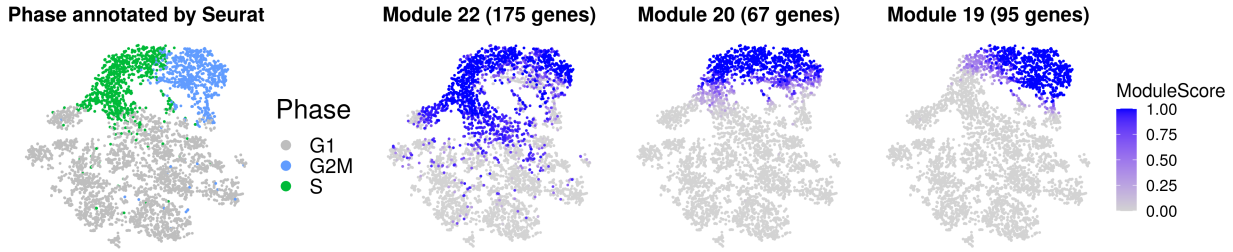


**Fig. S5.** Module scores of three cell cycle-associated gene modules identified from FACS-based mouse bone marrow dataset. Cell cycle annotations obtained from Seurat are shown on the left for reference. The t-SNE embedding is based on cell cycle marker genes (see Section 4.3).

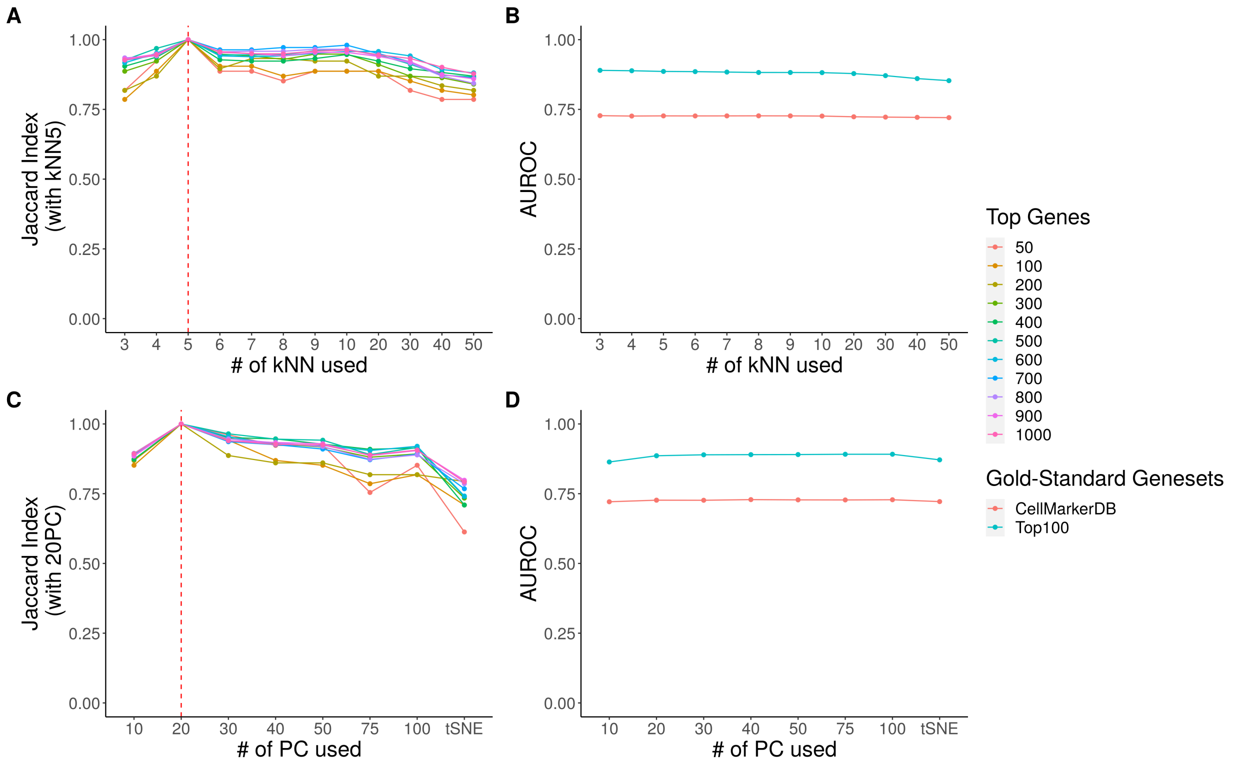


**Fig. S6.** LMD is robust to different numbers of neighbors $k$ and input spaces in constructing the cell-cell affinity graph. (A-C) Rank Stability: The Jaccard index between LMD's top N candidate markers for (A) $k = 5$ and other $k$, and (C) $20PCs$ and other input spaces. (B-D) AUROC Evaluation: Consistency of LMD's performance across (B) varying $k$, (D) varying input spaces, color-coded by two gold-standard gene sets: genes from the CellMarker database and the top 100 genes with the maximum fold changes across cell types.

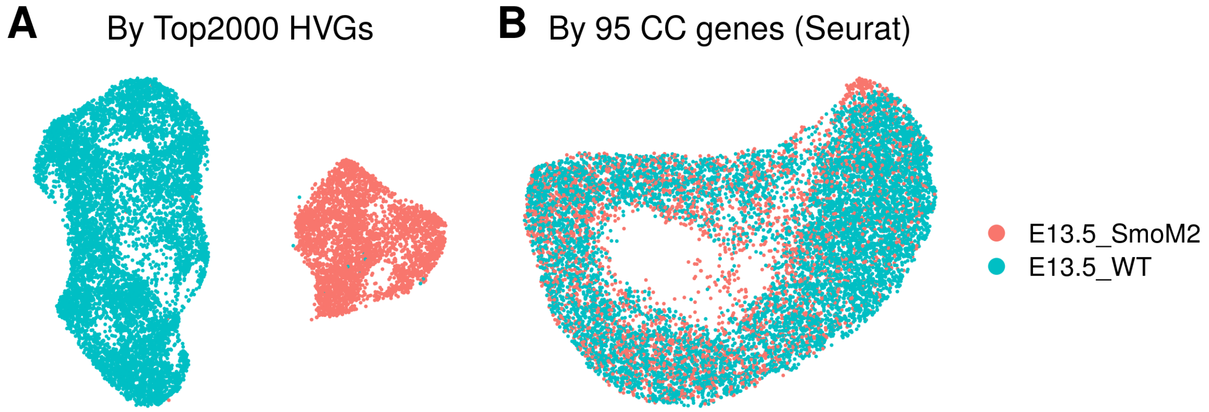


**Fig. S7.** Merge E13.5 SmoM2 and WT samples. (A) Embedding based on the top 2000 highly variable genes. (B) Embedding based on cell cycle marker genes (see Section 4.3).

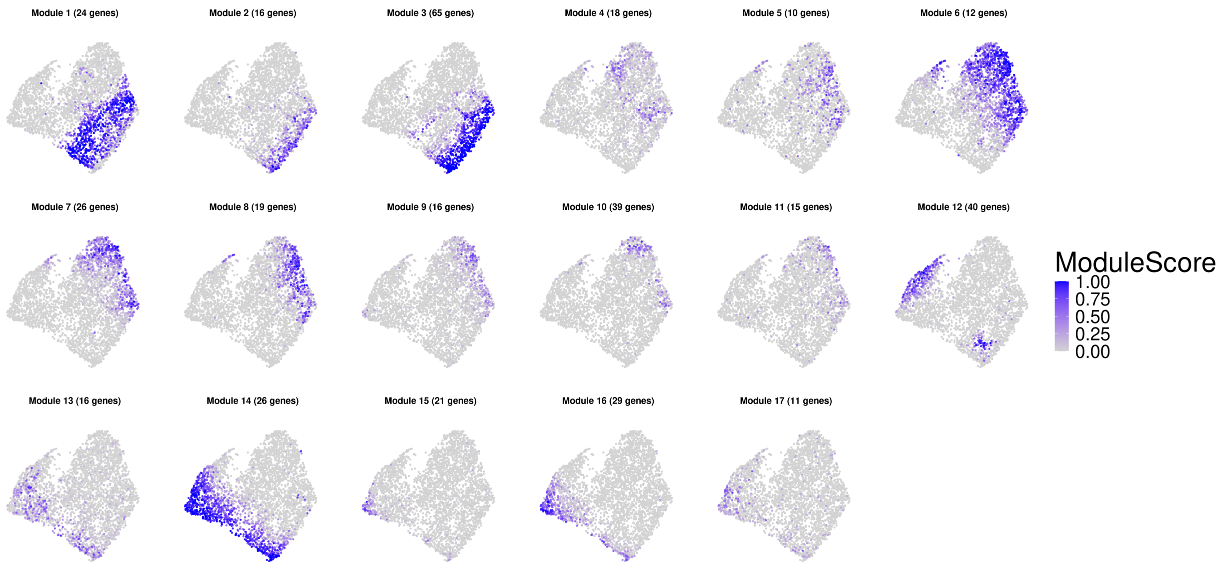


**Fig. S8.** UMAP of gene modules identified from E13.5 SmoM2 sample, color-coded by module score

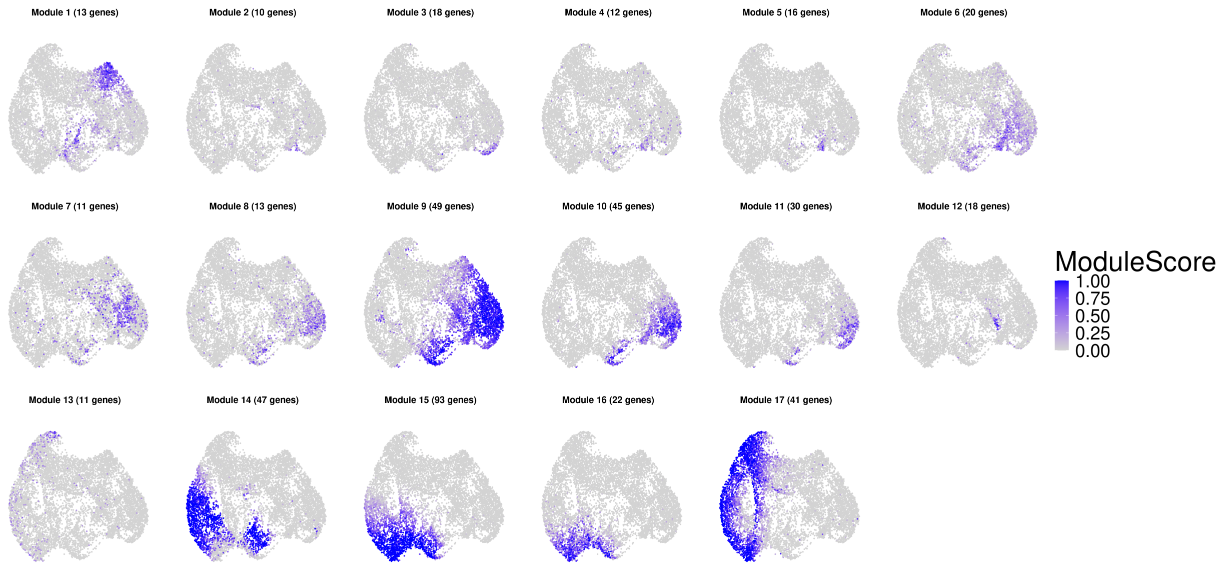


**Fig. S9.** UMAP of gene modules identified from E13.5 WT sample, color-coded by module score

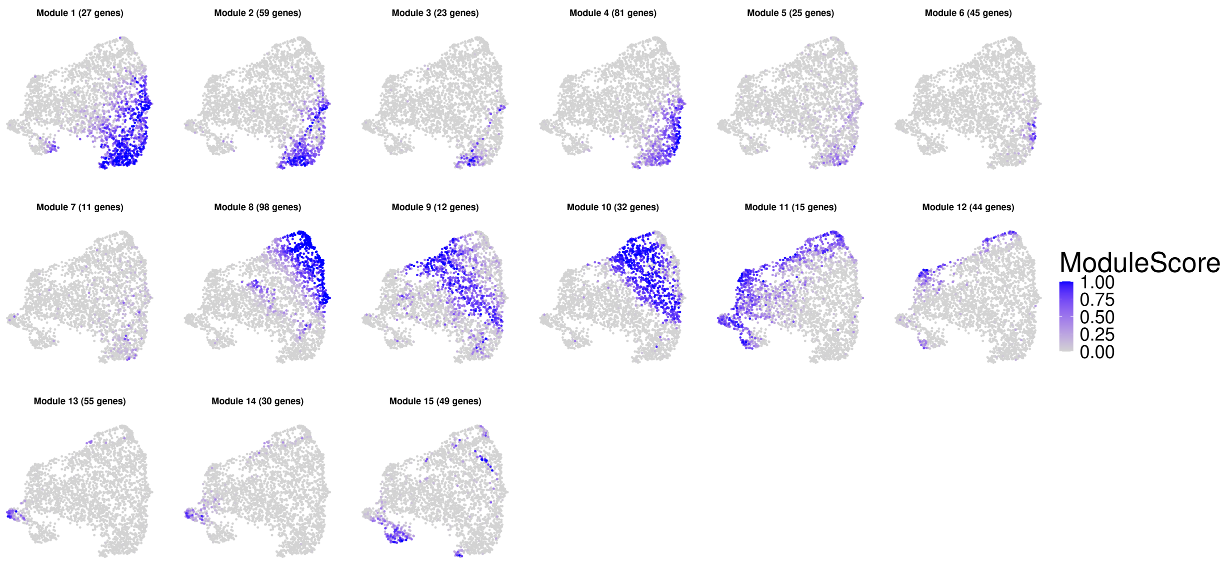


**Fig. S10.** UMAP of gene modules identified from E14.5 SmoM2 sample, color-coded by module score

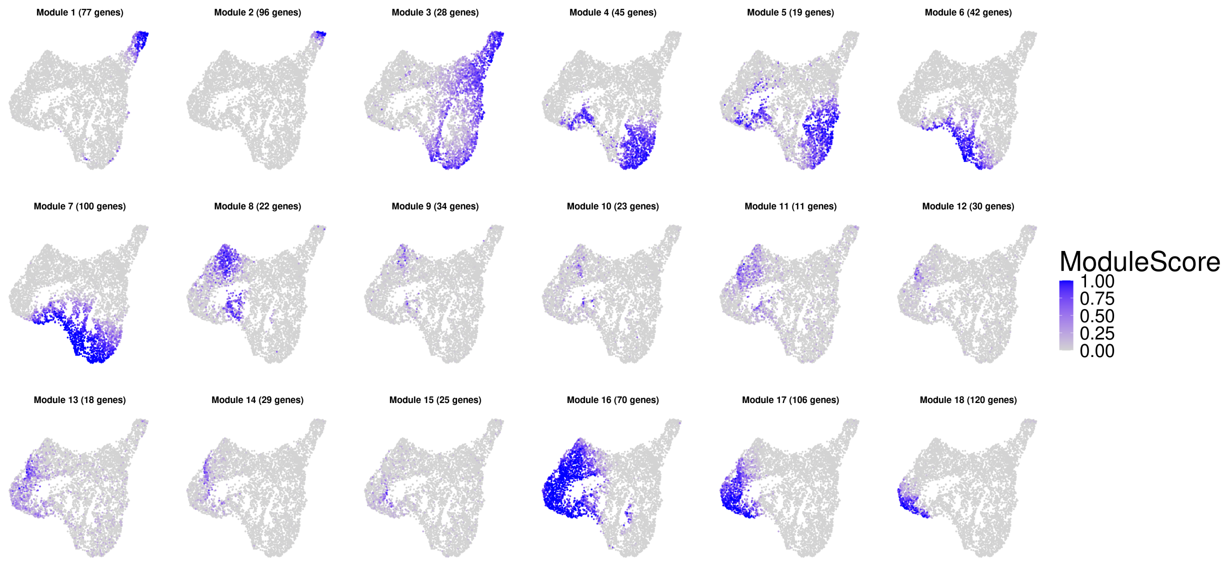


**Fig. S11.** UMAP of gene modules identified from E14.5 WT sample, color-coded by module score

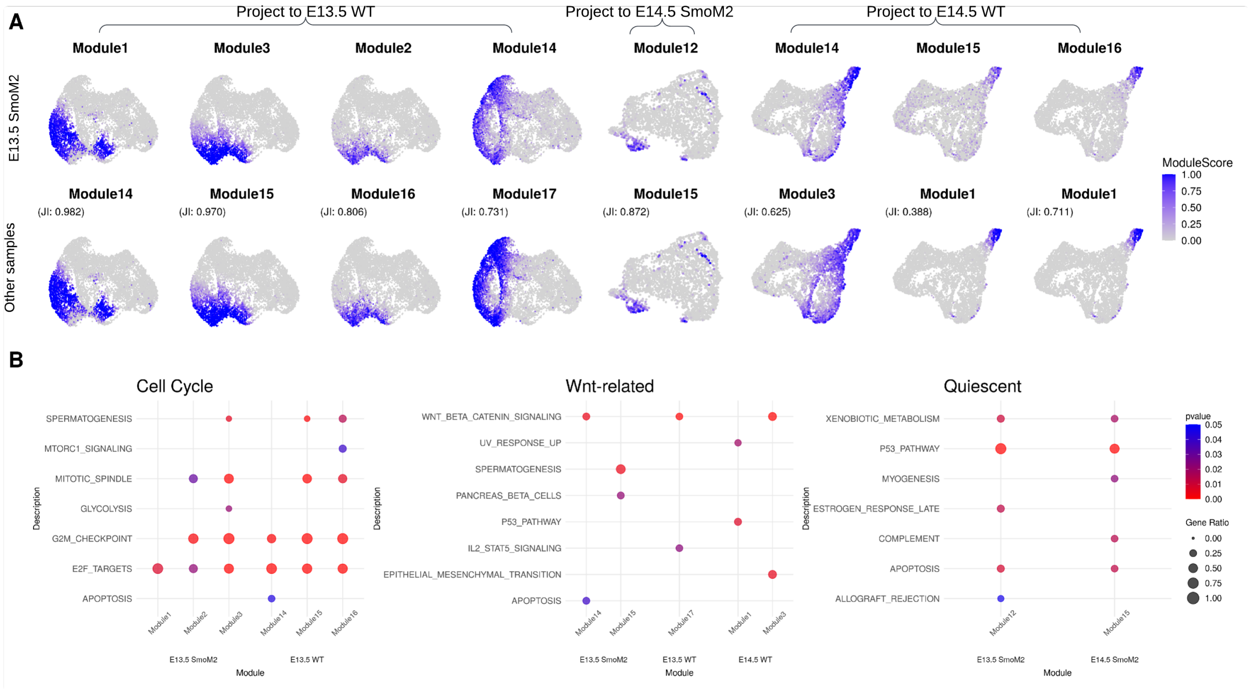


**Fig. S12.** Co-localization of E13.5 SmoM2 gene modules with gene modules from other samples. (A) Module scores of gene modules from E13.5 SmoM2 on other samples (first row) and gene modules derived from the corresponding samples with the highest association, determined by the Jaccard index (second row, as noted by the subtitle). (B) Bubble plot of the top 5 significantly enriched MsigDB pathways (pvalue$<$ 0.05) for each gene module listed above (non-significant pathways are excluded). Bubble color indicates p-value, and the size represents the fraction of module genes associated with the pathway.

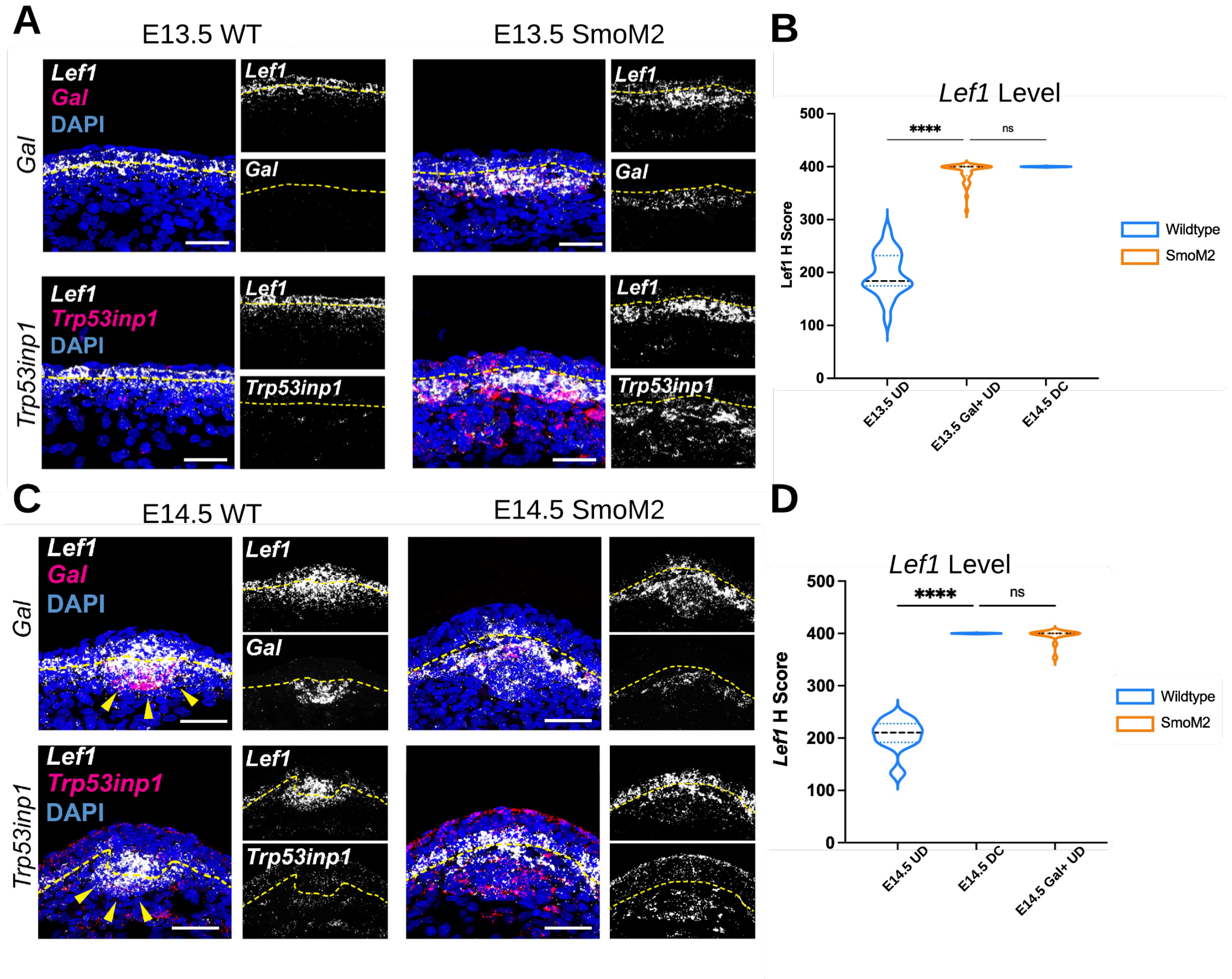


**Fig. S13.** Experimental validation of *Gal* and *Trp53inp1*, identified from module 16 and module 12 of E13.5 SmoM2, respectively. (A) FISH images (scale bar = 50 $\mu m$) showing the spatial distribution of *Gal* and *Trp53inp1* in the upper dermis of E13.5 WT and SmoM2, with *Lef1* as a comparison. (B) *Lef1* transcript levels (H-score) quantified by FISH in SmoM2 *Gal*+ cells at E13.5, compared to WT UDs at E13.5 and WT DCs at E14.5. (C) FISH images showing the spatial distribution of *Gal* and *Trp53inp1* transcripts in the upper dermis of E14.5 WT and SmoM2, with *Lef1* transcripts . (D) *Lef1* levels in SmoM2 *Gal*+ cells at E14.5, compared to WT UDs and DCs at E14.5.

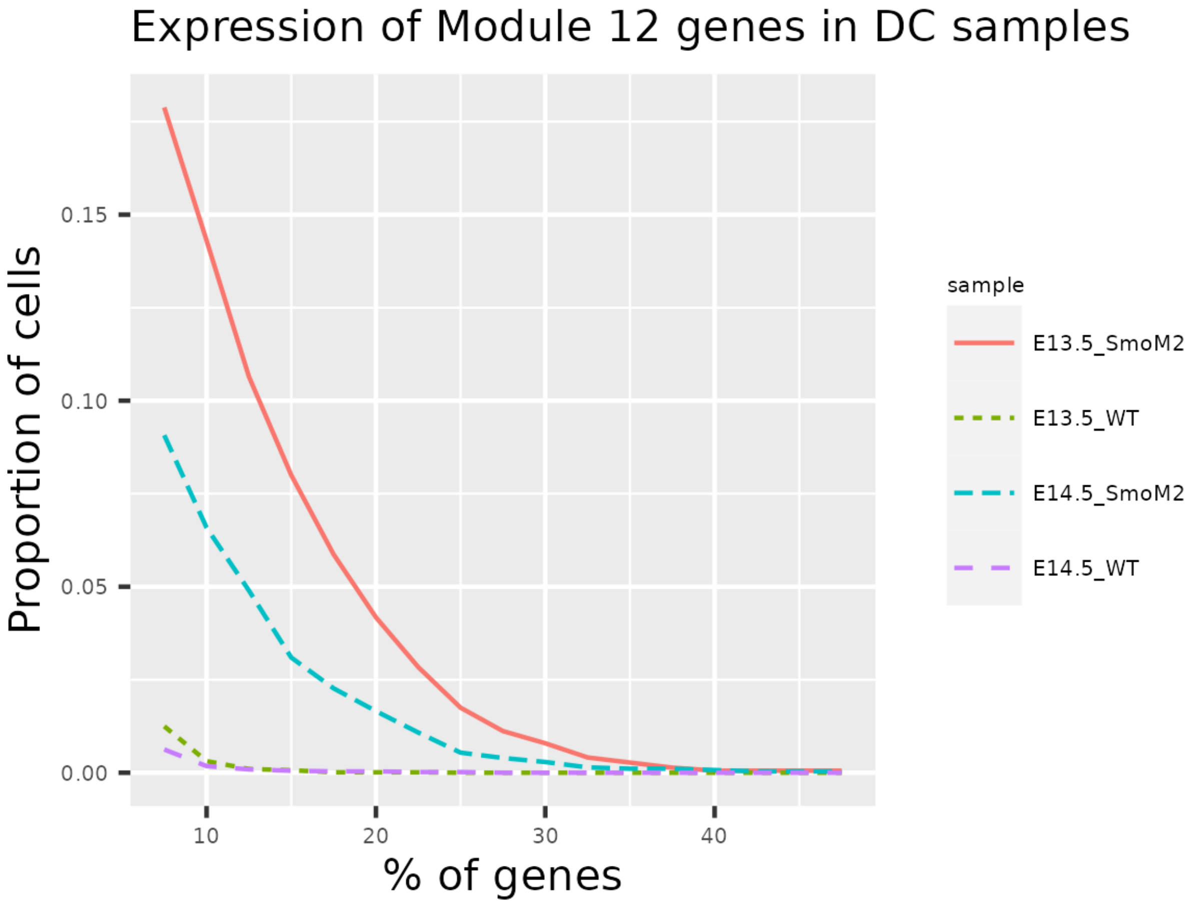


**Fig. S14.** Proportion of cells (y-axis) that expresses more than a percentage (x-axis) of genes from module 12 in each of the 4 hair follicle dermal condensates (DCs) samples.

### S2. Supplementary Tables

**Table S1.** The rank of selected genes in Fig. 2B across different methods

| | LMD | Seurat v4 | Hotspot | Marcopolo | SEMITONES | singlecellHaystack |
|---|---|---|---|---|---|---|
| Cd19 | 30 | 587 | 70 | 402 | 30 | 109 |
| Abca13 | 31 | 87 | 283 | 1142 | 54 | 474 |
| Il2rb | 75 | 14 | 214 | 251 | 1339 | 177 |
| Vpreb2 | 82 | 39 | 405 | 1398 | 5143 | 3335 |
| Mogat2 | 87 | 75 | 1087 | 1176 | 447 | 1764 |
| Phgdh | 501 | 1254 | 428 | 6 | 89 | 69 |
| Fyb | 539 | 1118 | 153 | 1651 | 215 | 39 |
| Grina | 2005 | 450 | 27 | 305 | 383 | 317 |
| Naaa | 1481 | 49 | 295 | 447 | 2479 | 2646 |
| Zf12 | 5147 | 11701 | 13123 | 1 | 11945 | 11367 |

**Table S2.** Runtime of LMD across various cell and gene numbers.

| nCells | nGenes | Time(minutes) |
|---|---|---|
| 1564 | 13108 | 0.52 |
| 1716 | 14348 | 0.62 |
| 2535 | 15649 | 0.67 |
| 5037 | 14585 | 2.49 |
| 6232 | 34446 | 5.04 |
| 5657 | 16284 | 8.34 |
| 10905 | 19301 | 9.13 |
| 21080 | 25949 | 26.19 |
| 30651 | 16074 | 43.57 |

### S3. Additional Resources

#### S3.1 Supplementary Excel

An Excel file (`Module_Description.xlsx`) containing detailed information of the 34 gene modules identified from Tabula Muris bone marrow FACS dataset, including module IDs, gene lists, associated cell types, and the top 5 most significantly enriched GO terms and pathways with BH adjusted p-values $< 0.05$, which can be downloaded here: `https://github.com/ruiqi0130/LMD_supplementary_data/raw/main/Module_Description.xlsx`

### Supplementary References

1. T. M. Consortium, O. coordination Schaum Nicholas 1 Karkanias Jim 2 Neff Norma F. 2 May Andrew P. 2 Quake Stephen R. quake@ stanford. edu 2 3 f Wyss-Coray Tony twc@ stanford. edu 4 5 6 g Darmanis Spyros spyros. darmanis@ czbiohub. org 2 h, L. coordination Batson Joshua 2 Botvinnik Olga 2 Chen Michelle B. 3 Chen Steven 2 Green Foad 2 Jones Robert C. 3 Maynard Ashley 2 Penland Lolita 2 Pisco Angela Oliveira 2 Sit Rene V. 2 Stanley Geoffrey M. 3 Webber James T. 2 Zanini Fabio 3, and C. data analysis Batson Joshua 2 Botvinnik Olga 2 Castro Paola 2 Croote Derek 3 Darmanis Spyros 2 DeRisi Joseph L. 2 27 Karkanias Jim 2 Pisco Angela Oliveira 2 Stanley Geoffrey M. 3 Webber James T. 2 Zanini Fabio 3, "Single-cell transcriptomics of 20 mouse organs creates a tabula muris," *Nature*, vol. 562, no. 7727, pp. 367–372, 2018.
